# The Mediator Kinase Module regulates cell cycle re-entry and transcriptional responses following DNA damage

**DOI:** 10.1101/2023.02.26.530133

**Authors:** Gönen Memişoğlu, Stefan Bohn, Nevan J. Krogan, James E. Haber, Alexander J. Ruthenburg

## Abstract

The Cdk8 kinase module (CKM) is a non-obligate and dissociable subcomplex of Mediator of transcription, a key regulator of RNA polymerase II (RNAPII). Through a genetic screen in yeast, we discovered a surprising role for Mediator CKM in the DNA damage response (DDR) and mitotic re-entry. Remarkably, we find that a single DNA break is sufficient for CKM-dependent global transcriptional attenuation. Upon DDR activation, the kinase activity of CKM antagonizes RNAPII binding to core Mediator, thereby reducing the transcriptionally-engaged RNAPII pool. This transcriptional attenuation is essential for DDR inactivation and limits the spreading of γ-H2AX into gene bodies. Furthermore, CKM localizes to DNA breaks to impede RNAPII binding. Importantly, we demonstrate that the role of CKM on DDR and transcriptional attenuation is conserved from yeast to mammals, establishing a multifaceted and essential function for CKM in transcriptional regulation of DNA-damage response.

## Introduction

To protect genomic integrity in the face of double strand breaks (DSBs), eukaryotic cells activate the DNA damage response (DDR)^1^, an elaborate and evolutionarily conserved signaling pathway orchestrated by the apical kinases Mec1/ATR andTel1/ATM^2^. In budding yeast, the DDR in response to a persistent DSB^3^ is driven predominantly by Mec1^4^ functioning through the downstream effector kinase Rad53/Chk2^5^, whereas Tel1 is largely dispensable^6^. Once recruited to DNA breaks via its binding partner Ddc2^7^, Mec1 phosphorylates histone H2A (referred to as γ-H2AX), which spreads extensively in *cis* around DSBs^8^ and in *trans* to nearby chromosomes^9^. About 12 hours after the induction of a DSB, Mec1 and Rad53 are inactivated^10^, which allows the cells to extinguish the DDR and re-enter mitosis^3^. The DDR, the kinases, as well as the kinase targets involved in this response, are highly conserved from yeast to mammals^11^. As in yeast, DSBs first stimulate ATM/Tel1 and ATR/Mec1, which then relay the signal to downstream effector kinases Chk1 and Chk2/Rad53 to enforce a cell cycle arrest^1^.

Mec1 and Rad53 are both essential for viability^12^, as they are required for replication fork stability under basal conditions^13,14^. The lethality of either *mec1*1 or *rad53*1 cells can be rescued by the upregulation of dNTP metabolism (e.g., by *sml1*1^15^). However, mounting genetic and biochemical evidence points to Mec1-independent roles for Rad53. For instance, Rad53 coordinates protein localization upon damage^16^, regulates histone levels^17^ and alters gene expression by directly binding to promoters^18^ independently of Mec1.

In addition to enforcing a mitotic arrest, Mec1 and Rad53 also modulate transcriptional responses to DNA damage^19,20^. The most extensively characterized aspect of this response is the Mec1- and Rad53-mediated upregulation of ribonucleotide reductase (*RNR)* genes to promote dNTP synthesis^21–24^. Surprisingly, beyond *RNR* genes and a few cell cycle-dependent transcripts, DNA damage does not alter the expression of genes essential for resistance to DNA damage^20,25,26^. Instead, DNA damage induced by ultraviolet radiation (UV) or replication stalling agent hydroxyurea (HU)^27^ triggers nonspecific downregulation of transcription via depletion of RNA polymerase II (RNAPII)^28–30^. This response is thought to prevent transcriptional stress resulting from RNAPII stalling at DNA lesions or transcription-replication fork collisions^31^. However, the impact of DNA damage that does not directly obstruct engaged RNA polymerase II (RNAPII) on transcription remains poorly characterized.

To investigate Mec1-independent functions of Rad53 and discover previously uncharacterized regulators of the DDR, we undertook a quantitative epistatic miniarray profile (E-MAP) screen^32,33^, which probes synthetic interactions between a query gene mutant and the yeast deletion library by systematic haploid crossing and selection. Our E-MAP analysis of *RAD53* and *MEC1* strains unveiled a genetic interaction between *RAD53* and several subunits of the Mediator of transcription. Mediator, an essential component of eukaryotic transcriptional machinery, bridges RNAPII to DNA-bound transcription factors and facilitates the assembly and firing of the preinitiation complex^34^. Core Mediator is composed of 21 highly conserved subunits alongside a handful of organism and preparation-specific factors grouped into head, middle and tail modules^35–38^. Core Mediator can also reversibly associate with an accessory Cdk8 Kinase Module (CKM)^39^, which consists of Cdk8 kinase, CycC, Med12 and Med13 in equimolar composition^40,41^. Structural and biochemical analyses indicate that the CKM-core Mediator and the RNAPII-core Mediator interactions are mutually exclusive^38,42–45^, accounting for observations of CKM as a general transcriptional repressor^46–48^. However, CKM can also *promote* transcription in a context-dependent manner^48–50^, especially in response to cellular stress^51–55^.

Here, after identifying a genetic interaction between *RAD53* and CKM in budding yeast, we explored the role of CKM in the context of the DDR. We find that CKM and its kinase activity are essential for mitotic re-entry following prolonged cell cycle arrest but are dispensable for DNA repair. Strikingly, a single DSB is sufficient to attenuate global transcription and this attenuation is a prerequisite for mitotic re-entry after damage. Akin to its repressor role under basal conditions, CKM mediates DNA damage-induced transcriptional repression by disrupting core Mediator-RNAPII interaction, thereby reducing RNAPII chromatin engagement in a kinase activity-dependent manner. Furthermore, we show that in the absence of CKM, the damage-specific histone mark γ-H2AX spreads into gene bodies. In addition to its global effect, CKM localizes to DNA breaks where it alters the DNA damage foci and antagonizes RNAPII recruitment. In the absence of CKM, RNAPII aberrantly binds to DSBs, driving mutagenic DNA repair. Finally, we demonstrate that CKM’s role in DDR regulation through transcriptional attenuation is conserved from yeast to mammals. CKM inhibition in human cell lines leads to aberrant DDR, elevated RNAPII chromatin retention, increased γ-H2AX foci and heightened sensitivity to DNA damaging reagents. Together, these findings establish CKM as a conserved and essential regulator of the DDR.

## Results

### E-MAP reveals a genetic interaction between *RAD53* and the Mediator kinase module

To discover factors that regulate the DNA damage response (DDR), we performed an E-MAP analysis^33^ of *RAD53 and MEC1*^56^ (Figure 1A, Table S1). In these assays, the lethality of *mec1*1 and *rad53*1 was suppressed by a *SML1* deletion^15^. As expected, under basal conditions, *RAD53* and *MEC1* phenotypic signature scores significantly overlap, especially for canonical DDR pathway components such as *RAD17*, *RAD24* and *RAD9*^57,58^ (Figure 1B). Surprisingly, the top scoring genetic interactions for *RAD53* are the subunits of the Mediator of transcription^34^ (Figure 1C). The genetic interactions between the CKM subunits *CDK8*, *CYCC*, *MED12* and *MED13* are specific to *RAD53* whereas the middle module subunits *MED1* and *MED9* display a strong correlation with both *RAD53* and *MEC1* (Figure 1B).

**Figure 1.**
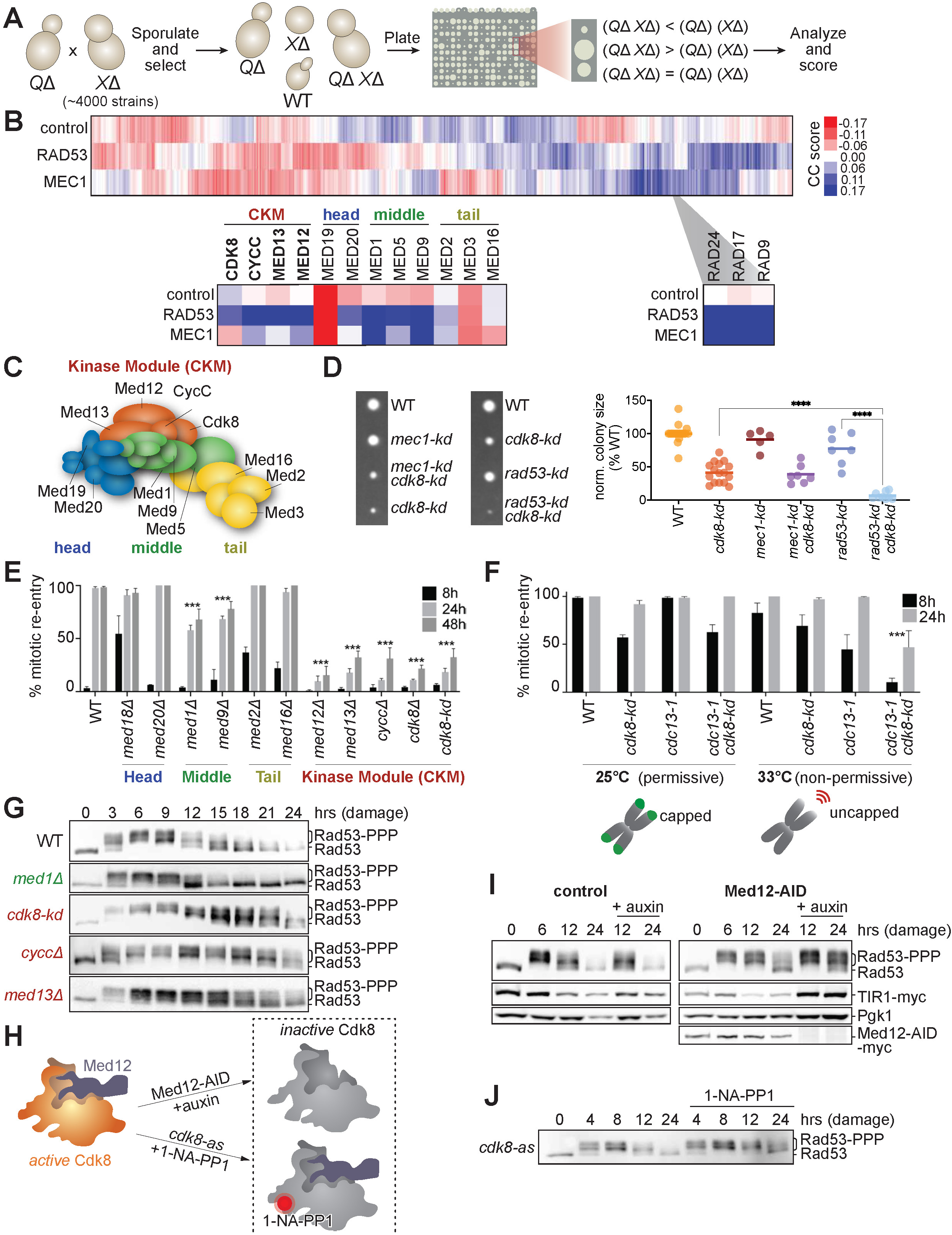
DDR kinase Rad53 and the kinase module of Mediator (CKM) synergistically impact growth and DNA damage response. **A.** Workflow of the epistatic miniarray profiling (E-MAP) screen^33^. Query strains (*Q*1) *rad53*1 *sml1*1 and *mec1*1 *sml1*1 were crossed to yeast deletion library strains (*X*1) to recover triple mutants. **B.** Heatmap representing the E-MAP scores for the control, *RAD53* and *MEC1*. **C.** Yeast Mediator complex, adapted from Guglielmi *et. al*^109^. **D.** Representative tetrads obtained from *mec1-kd cdk8-kd* and *rad53-kd cdk8-kd* diploids (left); and normalized colony sizes (right). All strains carry a compensatory *sml1*1 mutation (n>5; **** *p*<0.0001; one-way ANOVA). **E.** Percent mitotic re-entry following a DSB in Mediator mutants at indicated time points. **F.** Percent mitotic entry of *cdc13-1* strains with or without *cdk8-kd* mutation at permissive (25°C) or non-permissive (33°C) temperatures. **G.** Response to a DSB measured by Rad53 hyperphosphorylation in *med1*1, *cdk8-kd*, *cycc*1 and *med13*1. **H.** Two separate perturbation strategies (Med12-AID and *cdk8-as*) to conditionally inactivate CKM used in this study. **I-J.** Rad53 hyperphosphorylation following Med12-AID depletion (**I**), following the *cdk8-as* inactivation with 1-NA-PP1 (**J**).

To test whether the interaction between the CKM and *RAD53* is kinase activity-dependent, we crossed haploid *cdk8-kd* (kinase-dead)^59,60^ strains either with *rad53-kd*^61^ or with *mec1-kd*^62^. The *rad53-kd cdk8-kd* double mutant colonies were significantly smaller than WT or the single *kd* mutants (Figure 1D). The *rad53-kd cdk8-kd* double mutants exhibit increased histone expression compared to WT (Figure S1A), which partially accounts for the growth defect (Figure S1B), concordant with previous reports demonstrating that *rad53*1 accumulate excess histones which impede growth^17^. On the other hand, *mec1-kd cdk8-kd* double mutants only exhibited an additive reduction in growth, indicating that kinase activities of CKM and Rad53, not some other aspect of protein function.

### CKM mutations cause defects in the DNA damage response

To test whether CKM modulates the DDR, we created a collection of Mediator mutants in a strain background that contains a galactose-inducible HO endonuclease, which generates a single, irreparable DSB upon induction^3^. In WT cells, this DSB leads to a prolonged mitotic arrest in G_2_/M for about 12 hours, after which cells adapt and resume mitosis despite the persistent DSB. We quantified mitotic progression in these cells by microscopically tracking micromanipulated cells (Figure 1E). As expected, most WT cells arrested within 8 hours after DSB induction and resumed mitosis by 24 hours^3^. Mutants lacking head (*med18*1 and *med20*1) or tail (*med2*1 and *med16*1) Mediator subunits exhibited near-WT arrest and mitotic re-entry kinetics (Figure 1E). In contrast, the CKM mutants *med12*1*, med13*1*, cycc*1 and *cdk8*1, as well as the *cdk8-kd*, failed to re-enter mitosis at 24 hours following DNA damage. Middle module mutants *med1*1 and *med9*1, as well as the *med1*1 *med9*1 double mutant also showed mild mitotic re-entry defects (Figure 1E, S1C). These results are in accordance with structural analyses indicating that CKM binds to the core Mediator mainly via the Middle module^38,43^.

Next, we employed a *cdc13-1* mutant to generate DSBs without transcriptional activation of an endonuclease to test the mitotic re-entry defect of CKM mutants. In this strain, the *cdc13-1* mutation triggers telomere uncapping and DDR activation at nonpermissive temperatures^63^. While the *cdc13-1* cells arrested within 8 hours at nonpermissive temperature and re-entered mitosis by 24 hours, a significant portion of *cdc13-1 cdk8-kd* double mutants remained arrested at 24 hours (Figure 1F).

To complement these functional assays, we monitored the molecular readouts of DDR. Rad53 is hyperphosphorylated throughout the extended DSB-induced mitotic arrest and is gradually dephosphorylated as the cells adapt and resume mitosis^64^. In WT and *med1*1 cells, Rad53 phosphorylation is apparent 3 hours after a DSB, rising to apical levels after 6-8 hours and is fully dephosphorylated by 24 hours. However, in CKM mutants *med12*1, *med13*1 and *cdk8-kd*, Rad53 hyperphosphorylation persists at least for 24 hours, indicative of a mitotic re-entry defect (Figure 1G). None of the non-essential core Mediator mutants exhibited pronounced Rad53 phosphorylation defects in response to damage (Figure S1F), agreeing with our initial analyses (Figure 1E). These results demonstrate that CKM is required for a timely DDR inactivation and resumption of mitosis in response to a DSB.

Prolonged mitotic arrest in response to DSBs is maintained by DDR kinases Chk1 and Tel1, and the spindle-assembly protein Mad2^65–67^ (Figure S1E). We find that the mitotic re-entry defect of CKM mutants is dependent on Chk1 and Mad2 but is independent of Tel1 (Figure S1F). We note that *sml1*1, which suppresses the inviability of *mec1*1 and *rad53*1 E-MAP query strains, did not alter the mitotic arrest and re-entry of CKM mutants (Figure S1F). These results suggest that CKM modulates the Mec1-branch of the DDR signaling, even though it plays a Mec1-independent role under basal conditions.

To define the proximate role of CKM on DDR, we employed two independent conditional deactivation strategies (Figure 1H). First, we generated an auxin-inducible degron (AID)^68^ to deplete the CKM subunit Med12, which is essential for Cdk8 kinase activity^69,70^. Second, we created an analog-sensitive *cdk8-Y236G* mutant (hereafter referred to as *cdk8-as*), which is selectively inhibited by a large ATP analog 1-Naphthyl-PP1 (1-NA-PP1)^71^. Both acute Med12 depletion via auxin and *cdk8-as* inhibition via 1-NA-PP1 were sufficient to elicit an increased DDR upon DSB induction, evident from the accumulation of hyperphosphorylated Rad53 at 24 hours (Figure 1I, 1J, S1G). Therefore, DDR defects at both functional and molecular level are unique to the CKM and rely on its kinase activity.

### CKM mutants do not impact DNA repair

Next, we investigated the impact of CKM on DNA repair by homologous recombination via gene conversion (GC) or single strand annealing (SSA). In GC strain YJK17, a DSB on chromosome III is repaired by an ectopic donor on chromosome V via gene conversion within 8 hours^72^ without pronounced cell cycle arrest^73^ (Figure 2A). In SSA strain YMV80^64^, a DSB induced between two homologous sequences separated by 25 kb is repaired after extensive 5’ to 3’ resection (Figure 2E). This SSA repair requires approximately 12 hours to complete and induces cell cycle arrest until the repair is complete. In both GC and SSA strains, the viability of CKM mutants after damage was comparable to WT (Figure 2B, 2F), indicating that CKM mutants are proficient in homologous recombination by GC or SSA.

**Figure 2.**
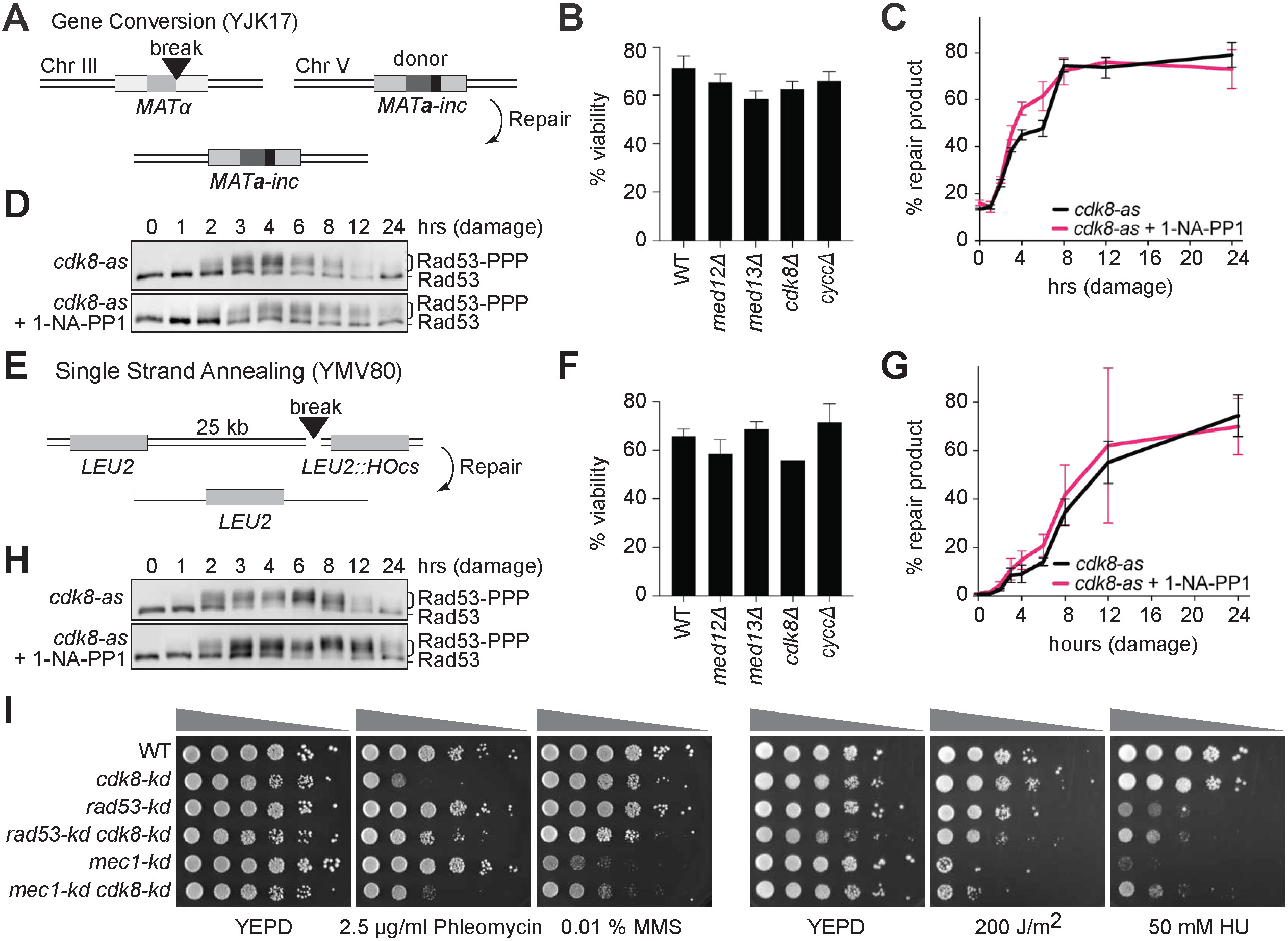
Loss of CKM does not alter DNA repair. **A.** In YJK17 gene conversion (GC) strain^72^, a DSB induced at *MAT*α (light gray) is repaired by using a *MAT***a**-inc ectopic donor (dark grey) that prevents continuous HO-endonuclease cutting, replacing, *MAT*α by *MAT***a**-*inc*. **B.** Viability of CKM mutants in YJK17 following damage. **C.** YJK17 repair kinetics, measured by primer extension assay^74^ in *cdk8-as* mutant with and without 1-NA-PP1 (n=3, error bars represent S.E.M.). **D.** DDR in YJK17 *cdk8-as* mutant with and without 1-NA-PP1 as measured by Rad53 hyperphosphorylation. **E.** In YMV80 single strand annealing (SSA) strain^64^, DSB between two homologous *LEU2* sequences (gray) leads to extensive end resection of the sequence between homologous sequences. The break is repaired when the homologous sequences exposed by resection anneal to each other. **F.** The viability of CKM mutants in YVM80 following damage. **G.** DNA repair kinetics measured by primer extension assay in YMV80 *cdk8-as* mutant with and without 1-NA-PP1 treatment (n=3, error bars represent S.E.M.). **H.** DDR in YMV80 *cdk8-as* mutant with and without 1-NA-PP1 as measured by Rad53 hyperphosphorylation. **I.** Sensitivity of *cdk8-kd*, *mec1-kd* and *rad53-kd* mutants to various DNA damaging drugs. All strains contain a *sml1*1 mutation.

Next, we measured repair kinetics by PCR^74^ in a *cdk8-as* mutant. Conditional CKM inhibition by 1-NA-PP1 treatment did not considerably alter repair kinetics or efficiency by GC (Figure 2C) or by SSA (Figure 2G). Lastly, we examined the DDR in these strains by Rad53 hyperphosphorylation following DNA damage. In the GC strain where the break is repaired rapidly, CKM inhibition did not cause a substantial difference in Rad53 hyperphosphorylation (Figure 2D). However, in the SSA strain, CKM inhibition resulted in a marked increase in Rad53 hyperphosphorylation (Figure 2H), indicating that CKM is dispensable for homologous recombination, but is essential for timely attenuation of DDR following successful DNA repair.

To further investigate the role of CKM in DNA repair, we assessed the sensitivity of *cdk8-kd* mutant to various DNA damaging drugs. The *cdk8-kd* mutant was not sensitive to low concentrations of replication stalling reagents methyl methanesulfonate (MMS) or hydroxyurea (HU) (Figure 2I), although it partially alleviated the HU sensitivity of *mec1-kd*^13,14^, suggesting that CKM loss does not cause replication stress. Although core Mediator has been implicated in nucleotide excision repair^75^, we find that *cdk8-kd* does not confer UV sensitivity, indicating that CKM is dispensable for this pathway. However, *cdk8-kd* is markedly sensitive to DSB-inducing reagent phleomycin; even more than *mec1-kd* or *rad53-kd* mutants. Given that CKM does not impair homologous recombination, the primary repair pathway in budding yeast, the phleomycin sensitivity of CKM mutants likely reflects a defect in mitotic re-entry, consistent with our previous analyses.

### CKM orchestrates to the global downregulation of transcription in response to DNA damage

Given the function of CKM in transcription^34^, we hypothesized that CKM could interface with DDR by regulating the transcriptional responses to DNA damage. To examine this, we performed RNA sequencing following acute inhibition of CKM activity by Med12-AID depletion (Figure S2A, B). Consistent with previous reports^42,76^, CKM depletion alone caused a modest increase in global transcription even after 10 hours of CKM depletion (Figure 3A, B). In contrast, global transcription was substantially attenuated after a single DSB. Remarkably, this downregulation is largely dependent on CKM, as CKM inactivation reversed the transcriptional attenuation of nearly all transcripts (Figure 3C), including transcripts involved in DDR, cell cycle and dNTP metabolism (Figure S2D). We note that the transcriptional response to a DSB differs from the transcriptional responses to HU^18^ or MMS^19^ (Figure S2E, S2F) and is not limited to silencing of DSB-proximal genes due to DNA end-resection^25,77^. These data suggest that CKM plays a prominent and distinct role in transcriptional attenuation triggered by DSBs as compared to its modest impact on transcription under basal conditions.

**Figure 3.**
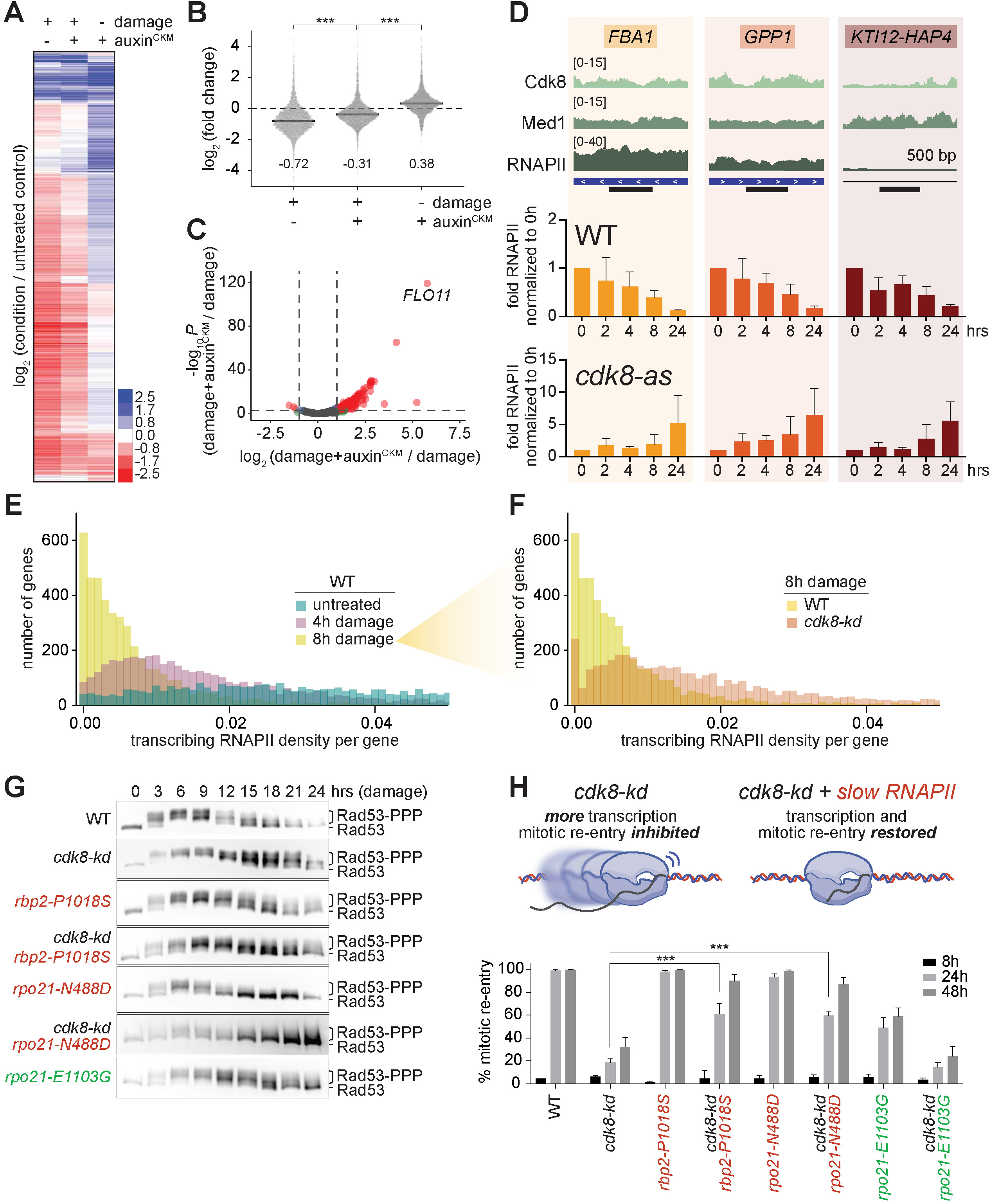
A single DNA break is sufficient for global shut-down of transcription and global displacement of RNAPII from chromatin. **A-B.** Differential expression analysis from RNA sequencing of Med12-AID cells (n=3 per condition), after a DSB (8 hours) and CKM depletion by auxin treatment (10 hours), represented as (**A**) a heatmap of normalized log_2_-fold change, (**B**) an aggregate plot (gray bars represent median, \*\*\**p*<0.0001, one-way ANOVA). **C.** Volcano plot of CKM-dependent changes in expression following a DSB (*p*-value cutoff=10^-4^, fold-change cutoff=1). **D.** (Above) Normalized Cdk8, RNAPII (Rbp3) and core Mediator (Med1) enrichment in untreated cells from Rossi *et al*.^78^. Three control loci (*FBA1*, *GPP1* and *KTI12-HAP4*) were picked based on their varying level of RNAPII enrichment for further ChIP-qPCR analysis. (Below) normalized fold RNAPII (Rbp3-V5) enrichment at the three control loci in a WT strain and a *cdk8-as* strain treated with 1-NA-PP1 following a DSB, assayed by ChIP-qPCR (n=3, error bars represent S.E.M., qPCR amplicon depicted as a black bar below genome tracks). **E-F.** Transcribing RNAPII density per gene as measured by PRO-seq following a DSB in (**E**) WT cells (**F**) in *cdk8-kd* cells. Strand-specific PRO-seq read counts were normalized to gene length and plotted as a histogram (X-axis truncated for clarity, see Figure S4A for extended histograms). **G.** DDR to a single DSB in fast (*rpo21-E1103G*; green) and slow (*rbp2-P1018S* or *rpo21-N488D*; red) RNAPII mutants measured by Rad53 hyperphosphorylation. **H.** Percent mitotic re-entry in fast and slow RNAPII mutants.

In response to DNA damage by UV, cells suppress transcription by ubiquitylation and subsequent degradation of RNAPII^30^. To address whether a DSB downregulates transcription through the same mechanism, we created an RNAPII (*rpo21-K1246R*) mutant which cannot be ubiquitylated and therefore is stabilized. However, we did not detect any cell cycle arrest and re-entry defects in this mutant (Figure S2G). Moreover, we do not observe notable changes in abundance RNAPII subunits Rbp3 and Rbp1 or the core Mediator subunit Med1 following double-strand breaks (Figure S2H). Additionally, the inhibition of CKM activity in *cdk8-as* (Figure S2H) or via Med12 depletion (Figure S2C) did not alter the abundance of RNAPII. However, we do detect a steady decrease in Cdk8 levels upon inhibition of its kinase activity (Figure S2H, discussed below). Therefore, the drastic transcriptomic repression in response to a DSB cannot be attributed to the depletion of core transcriptional machinery; rather, CKM plays a central role in this attenuation.

### CKM limits chromatin-bound RNAPII pools to inhibit transcription upon DNA damage

We posited that the DNA damage-dependent transcriptional attenuation might stem from a reduction in RNAPII chromatin occupancy. To test this, we chromatin immunoprecipitated (ChIP) RNAPII subunit Rbp3 at three control loci far from the DSB site, that display a range of basal of Cdk8, Med1 and RNAPII enrichment (Figure 3D)^78^. In WT cells, RNAPII/Rbp3 chromatin occupancy steadily decreased at all three loci after a DSB, independent of gene length and basal CKM, core Mediator and RNAPII occupancy levels (Figure 3D), consistent with global transcriptional attenuation upon damage. This damage-dependent decrease in RNAPII chromatin occupancy was completely reversed by CKM inhibition in the *cdk8-as* mutant with 1-NA-PP1. We note that neither DNA damage nor CKM depletion led to extreme transcriptional changes at these control loci (Figure S2I).

As ChIP does not distinguish between inactive chromatin-bound RNAPII and transcriptionally engaged RNAPII, and RNA-seq is an indirect measure of transcriptional activity, we employed precision run-on sequencing (PRO-seq)^79,80^ to directly measure nascent transcription. PRO-seq revealed that actively transcribing RNAPII pools drastically decrease in WT cells following a DSB (Figure 3E, Figure S3A). In accord with the ChIP results, this damage-dependent reduction in transcribing RNAPII was also dependent on CKM, as *cdk8-kd* cells maintained increased levels of active RNAPII relative to WT upon damage (Figure 3F, S3A). These findings indicate that CKM attenuates global transcription in response to damage by antagonizing active RNAPII.

### Transcriptional downregulation is essential for mitotic re-entry following DNA damage

Next, we asked whether transcriptional downregulation is a prerequisite for mitotic re-entry after DNA damage. To investigate this, we introduced mutations in RNAPII which either increase (*rpo21-E1103G*; referred to as *fast*) or decrease (*rbp2-P1018S* and *rpo21-N488D*; referred to as *slow*) RNAPII elongation rate and transcriptional output^81^. When challenged with a DSB, the slow RNAPII mutants re-entered mitosis much earlier than WT, evident in the Rad53 hyperphosphorylation analysis of *rpo21-N488D* (Figure 3G). Conversely, the fast RNAPII mutant failed to re-enter mitosis by 24 hours after the induction of a DSB, apparent from the single-cell microscopy (Figure 3H) and the Rad53 hyperphosphorylation assays (Figure 3G). Strikingly, downregulating transcription in *cdk8-kd* mutant by introducing slow RNAPII mutations rescued the mitotic re-entry defect and prevented accumulation of hyperphosphorylated Rad53 species at 24 hours (Figure 3G, 3H). Combining RNAPII fast mutant *rpo21-E1103G* with *cdk8-kd* did not significantly worsen the defect of *cdk8-kd*, indicating that CKM is the main antagonist of RNAPII in response to damage and that this antagonism is critical for cell cycle re-entry. Furthermore, *cdk8-kd* mutation marginally rescued the phleomycin sensitivity of slow mutants, whereas the *cdk8-kd rpo21-E1103G* (fast) double mutant was more sensitive to phleomycin than the single mutants (Figure S3B).

To validate these findings, we inhibited transcription by auxin-depleting the essential core Mediator subunit Med17^82^. In WT cells, Med17 depletion did not alter Rad53 hyperphosphorylation kinetics after damage (Figure S3C). However, Med17 depletion rescued the mitotic re-entry defect of *cdk8-kd* cells, evident from the reduced Rad53 hyperphosphorylation levels. In contrast, Med17 depletion was not sufficient to rescue the mitotic re-entry defect of *ptc2*1, which is caused by a failure of Rad53 inactivation^83^. These data suggest that mitotic re-entry defect of CKM mutants is due to elevated transcription during the DDR, and increased transcriptional output is detrimental for mitotic re-entry following DNA damage.

### CKM kinase is active and intact during DNA damage response

We posited that DNA damage could alter CKM’s kinase activity to initiate damage-dependent transcriptional inhibition. To test this, we utilized Med2, a previously characterized CKM target, as a read-out of CKM activity^84^. In WT cells, phosphorylated Med2 results in a hypomobility shift on a western blot (Figure S4A). This shift was absent in a *med2-S208A* mutant, which abolishes the CKM target site, and was reduced in a *cdk8-kd* strain, confirming prior results^84,85^ (Figure 4A). We note that *med2-S208A* mutation did not affect DDR as assayed by Rad53 hyperphosphorylation (Figure S4B). We detect an apparent increase in Med2 phosphorylation by this assay in our WT strain 4 hours after a DSB, which gradually reverts to the phosphorylation levels observed in *cdk8-kd* mutants (Figure 4A). This Med2 phosphorylation is also evident in WT strains 8 hours after nocodazole treatment which causes a G_2_/M arrest^86^ (Figure 4B). Furthermore, Med2 phosphorylation was depleted upon CKM inhibition in *cdk8-as* cells treated with 1-NA-PP1 (Figure 4C). Albeit indirectly, these results indicate that CKM kinase activity is stimulated upon G_2_/M arrest induced by DNA damage and reduced as cells resume mitosis.

**Figure 4.**
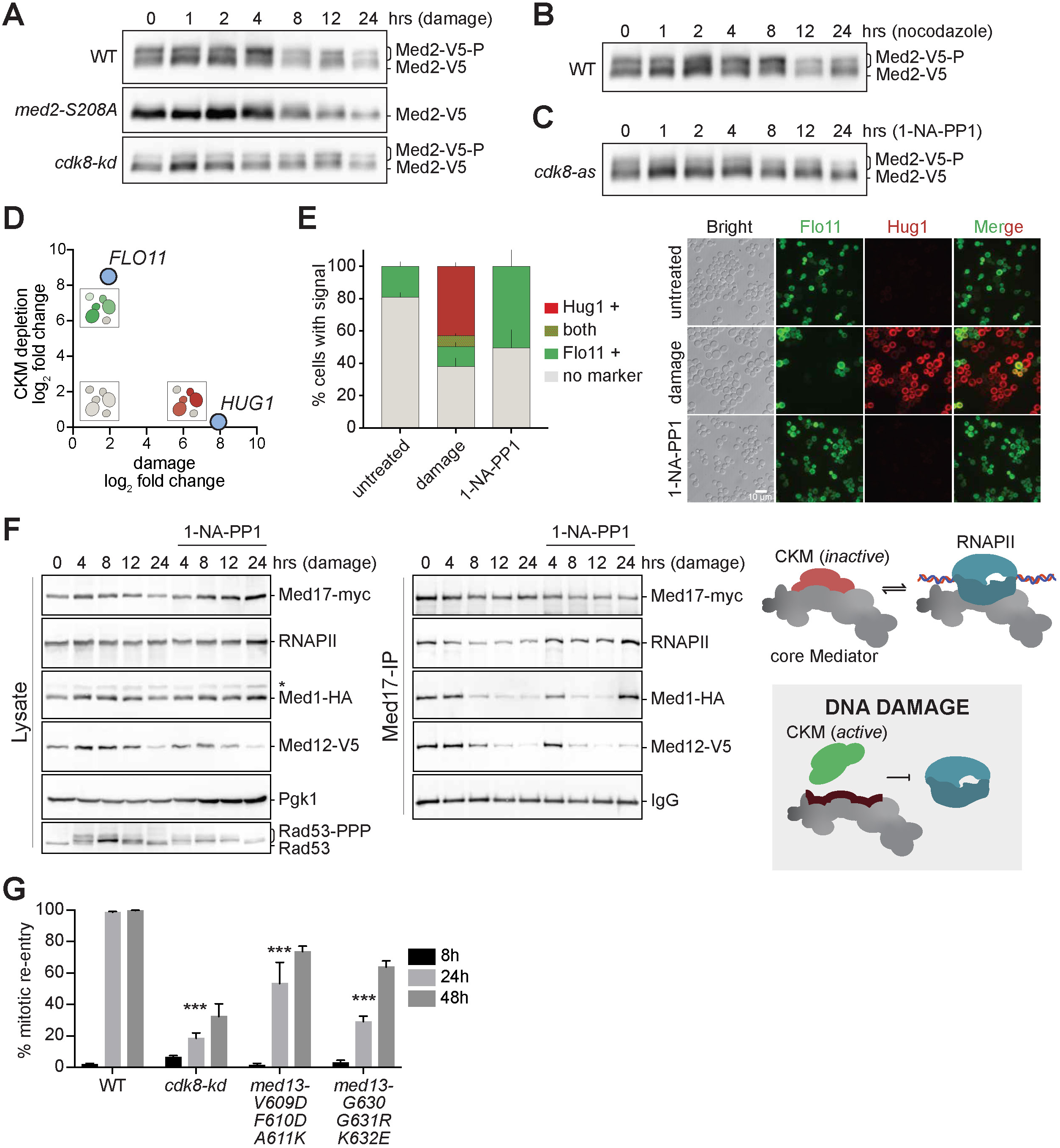
Kinetically active CKM antagonizes core Mediator and RNAPII binding in response to DNA damage. **A.** Med2 phosphorylation in WT, *med2-S208A* and *cdk8-kd*, detected as a hypomobility shift with a Western blot. **B.** Med2 phosphorylation in WT following nocodazole treatment. **C.** Med2 phosphorylation in *cdk8-as* with 10 µM 1-NA-PP1. **D.** Changes in CKM-responsive transcript *FLO11* and damage-responsive transcript *HUG1* after CKM depletion versus DNA damage, extracted from differential expression analysis (Figure 3A). **E.** (right) Number of cells with detectible levels of Flo11::mNeonGreen or Hug1::mScarletI in *cdk8-as*, following 8 hours damage or 1-NA-PP1 treatment; (right) representative microscopy images of *cdk8-as* Flo11::mNeonGreen or Hug1::mScarletI cells. **F.** Core Mediator (Med17-myc and Med1-HA), RNAPII and CKM (Med12-V5) interactions following damage in a *cdk8-as* strain, with or without 1-NA-PP1. Samples collected at various time points were assayed by Western blotting after Med17-myc coimmunoprecipitation (right). Cell lysate samples collected from the same cultures (left) were blotted for Rad53 to infer DDR and for Pgk1 as loading control. **G.** Percent mitotic re-entry of Med13 mutants (*V609D/F610K/A611K* and *G630R/G631R/K632E*).

Our expression analyses (Figure 3A) reveal that *FLO11* transcript abundance is inversely correlated to CKM activity, whereas *HUG1* abundance is directly correlated to DDR (Figure 4D), as shown previously^24^. To test the activity of CKM following DDR activation on a single cell level, we leveraged the orthogonal specificity of these transcripts to CKM depletion and DNA damage respectively, and generated a reporter strain in which Flo11 and Hug1 proteins are fluorescently labeled. None of the WT cells expressed these fluorescent reporters under basal conditions (Figure S4C), highlighting the specificity of these reporters to the respective stimuli. However, a small percentage of untreated *cdk8-as* cells expressed Flo11, indicating that this analog-sensitive mutation leads to a mild loss of CKM function (Figure 4E). Regardless, the percentage of Flo11 expressing *cdk8-as* cells increases significantly upon CKM inhibition via 1-NA-PP1 treatment as expected. DNA damage led to increased Hug1 expression, whereas the fraction of Flo11-expressing cells remained comparable to the levels observed in the untreated control (Figure 4E), indicating that CKM is kinetically active while DDR is active, consistent with our Med2 phosphorylation results (Figure 4B).

To monitor DNA damage-dependent changes to CKM stoichiometry and subunit abundance, we appended all CKM subunits with epitope tags in a single strain (Figure S4D). We observe that initially, a DSB leads to a uniform upregulation of all four CKM subunits, which is consistent with biochemical and structural studies that define CKM as equimolar in subunit composition^37,41,43^. This upregulation is followed by a steady decrease in CKM subunit abundance at 24h both in cycling and DNA damage-treated cultures, which is consistent across several experiments (Figure 1I, S2C, S4D, 4F, S4F). Notably, the changes in CKM abundance upon damage correlate with the changes in CKM activity assayed via Med2 phosphorylation, implying an autoregulatory mechanism that modulates CKM abundance. Our analyses of CKM stoichiometry by Cdk8-FLAG and Med13-myc immunoprecipitation (Figure S4E) also indicate that DNA damage does not significantly alter the composition of the complex. Collectively, these results imply that DDR alters CKM activity and abundance, but not the subunit stoichiometry, to modulate transcription.

### CKM activity is required to inhibit core Mediator binding to RNAPII after DNA damage

We find that nascent transcription is attenuated and RNAPII chromatin recruitment is reduced in response to a DSB. To test whether DNA damage suppresses transcription initiation, we analyzed the damage-dependent changes to preinitiation complex assembly. We coimmunoprecipitated the essential core Mediator subunit of Med17, which directly interacts with RNAPII in the preinitiation complex^44^, in a strain containing epitope-tagged core Mediator subunit Med1 and CKM subunit Med12 (Figure 4F, Figure S4F). We find that Med17-RNAPII as well as Med1-Med17 interactions are gradually reduced following a DSB (Figure 4F). We note that apparent Med17-RNAPII binding is not mediated by DNA or RNA, as benzonase digestion did not alter this interaction (Figure S4G). Given that core Mediator-RNAPII binding is indispensable for transcription initiation, these findings hint that transcription initiation is suppressed following DNA damage.

Our results indicate that inhibition of CKM kinase activity is sufficient to counteract the damage-dependent transcriptional attenuation. To test whether CKM kinase activity alters preinitiation complex in response to damage, we performed the same coimmunoprecipitation experiments with *cdk8-as* cells in the presence of 1-NA-PP1 to inhibit CKM activity. CKM inhibition prevented the damage-dependent dissociation of RNAPII from core Mediator subunit Med17, consistent with a defect in transcription initiation. Interestingly, upon CKM inhibition, Med1-Med17 and RNAPII-Med17 binding are enriched specifically at 24 hours after damage, which is also reflected in our RNAPII-ChIP analysis, showing increased RNAPII chromatin binding (Figure 3D). Collectively these data indicate that upon damage CKM antagonizes core Mediator-RNAPII interaction in a kinase activity-dependent manner. Reduced core Mediator-RNAPII interaction, in turn, leads to reduced transcriptional output, consistent with our RNA-seq, ChIP-qPCR and PRO-seq results.

Structural and biochemical studies demonstrate that CKM and RNAPII compete for the same binding interface on core Mediator^38,42–45^. Surprisingly, we find that the steady decrease in CKM abundance following DNA damage does not translate to increased core Mediator-RNAPII binding, revealing that CKM counteracts core Mediator-RNAPII binding upon DNA damage via a kinase-dependent mechanism instead of steric obstruction. Corroborating this, inhibition of CKM kinase activity is sufficient to upregulate core Mediator-RNAPII binding (Figure 4F). To further validate the results from these binding assays indicating a direct role for CKM in antagonizing core Mediator-RNAPII binding, we mutated the core Mediator binding interface of CKM (Med13), which impairs CKM-core Mediator binding^38^. These core binding-defective Med13 mutants significantly reduced mitotic re-entry following DNA damage, mimicking a loss of CKM (Figure 4G). Based on these results, we postulate that upon damage, CKM binds to core Mediator and alters its binding interface that is shared between CKM and RNAPII likely through post-translational modifications, insulating the core Mediator away from RNAPII to inhibit transcription initiation.

### CKM functions locally at DNA break sites

DSBs inhibit transcription of break-proximal genes both in mammals^87^ and in yeast^25,77^. To test whether CKM contributes to this transcriptional repression near DSBs, we assayed the localization of core Mediator (Med1), CKM (Cdk8) and RNAPII (Rbp3) around two separate DSBs by ChIP-qPCR (Figure S2F). In WT cells upon damage, Med1 and Cdk8 localized to DSBs, while RNAPII was excluded (Figure 5A, Figure S5A). CKM inhibition in *cdk8-as* mutant blocked the damage-dependent Cdk8 and Med1 binding to DSBs; while RNAPII recruitment to DSBs was elevated (Figure 5A, right). The enrichment of RNAPII at DSBs after CKM inhibition was highest at 24 hours, analogous to RNAPII enrichment results for the three control loci (Figure 3D). These findings suggest that CKM functions both locally at DSBs and globally to restrict RNAPII chromatin recruitment to antagonize transcription.

**Figure 5.**
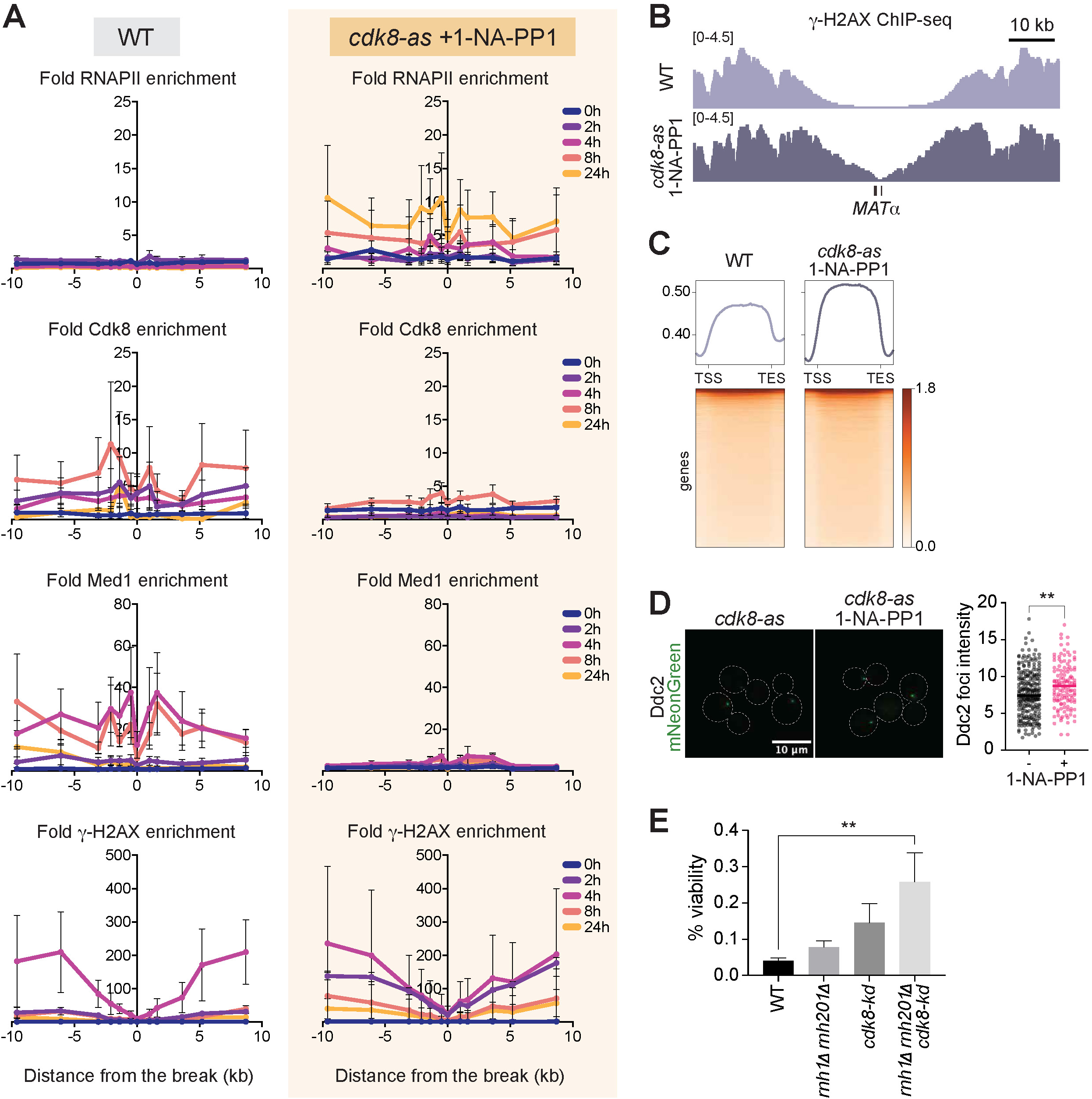
CKM prevents RNAPII recruitment to DNA breaks. **A.** Fold enrichment of RNAPII (Rbp3-V5), Cdk8 (Cdk8-myc), core Mediator (Med1-HA) and γ-H2AX around DSB measured by ChIP-qPCR in a WT and *cdk8-as* strain treated with 1-NA-PP1 (n>2, error bars represent S.E.M.). **B.** ChIP-seq coverage of γ-H2AX around the DSB at 8 hours. **C.** Aggregate plot of normalized γ-H2AX ChIP signal around transcription start sites (TSS), 8 hours after DSB. **D.** Viability in WT, *cdk8-kd*, *rnh11 rnh2011* mutant cells (n>3; error bars represent S.E.M., one-way ANOVA, \*\**p*=0.0053). **E.** Ddc2 foci intensity 8 hours after damage in *cdk8-as* cells, with or without 1-NA-PP1 (Welch’s t-test, \*\**p*=0.017).

To test whether CKM activity at the break site impacts local features of DDR, we analyzed the enrichment of γ-H2AX, a DNA damage-dependent histone mark which rapidly spreads around DSBs^8^. CKM inhibition did not alter the γ-H2AX enrichment around DSBs, measured by ChIP-qPCR (Figure 5A); however, it did increase break-proximal γ-H2AX spreading, measured by γ-H2AX ChIP-seq (Figure 5B, Figure S5B). In addition to DSBs, γ-H2AX spreads to nearby chromosomes in *trans*^9^ and the enrichment of γ-H2AX is inversely correlated with transcriptional activity^9,88^. Given CKM’s critical role in regulating global transcription upon DNA damage, we posited that it could alter global γ-H2AX spreading. Our aggregate analysis of γ-H2AX ChIP-seq signal 8 hours after damage shows that CKM inhibition did not alter markedly γ-H2AX spreading at transcriptional start sites (TSSs) (Figure S5C); however, it resulted elevated γ-H2AX enrichment within RNAPII-dependent gene bodies (Figure SC). These findings indicate that CKM-dependent transcriptional attenuation could also restrict γ-H2AX spreading into transcriptionally active sites following DNA damage.

Mec1 phosphorylates γ-H2AX upon localization to DSBs via its binding partner Ddc2^7,89^. Ddc2 focus intensity is correlated with the Mec1 kinase activity^10^. We asked if in CKM-dependent changes at DSBs could alter Ddc2-Mec1 recruitment, which in turn would alter DDR and γ-H2AX spreading. Our analysis reveals that CKM-inhibited cells display enhanced Ddc2-mNeonGreen focus intensity at 8 hours after damage (Figure 5D), suggesting that CKM-dependent changes to DNA damage foci modify the recruitment or stabilization of DDR machinery to DSBs, which results in elevated γ-H2AX spreading.

DSBs near transcriptionally active regions can give rise to translocations via mutagenic repair pathways such as end-joining^90,91^. We postulated that CKM-mediated exclusion of RNAPII from DSBs could prevent such mutagenic end-joining events. To test this, we assayed the viability of strains that induce a single and irreparable DSB at the *MAT*α locus and repair via nonhomologous end-joining^92^. The viability of *cdk8-kd* after damage was modestly increased compared to WT cells, although this increase was not statistically significant (Figure 5E). To further stabilize the labile transcripts from active transcription at DSBs that could participate in DNA repair^93^, we deleted the RNase H1 (*RNH1*) and H2 (*RNH201*). The viability of *cdk8-kd rnh1*1 *rnh201*1 triple mutant was nearly 5-fold higher than the WT (Figure 5E). Therefore, in addition to its global role in transcriptional downregulation, CKM acts locally to limit the localization of transcriptional complexes to DSBs, which could result in pervasive transcription and mutagenic DNA repair.

### The function of CKM is conserved from yeast to mammalian cells

Although mammalian Mediator contains additional subunits and paralogs, the structure and the main function of the complex is highly conserved from yeast to mammals^47^. To examine the role of CKM in mammalian DDR, we employed the U2OS cell line^88^; which contains the restriction enzyme *Asi*SI fused to the estrogen receptor (ER) hormone-binding domain. Upon treatment with 4-hydroxytamoxifen (4-OHT), *Asi*SI-ER translocates to the nucleus, inducing nearly 170 DSBs^94^. The 4-OHT-dependent DSB induction led to the activation of DDR kinases ATM, DNA-PKcs, Chk1 and Chk2 within 4 hours, without altering transcriptional machinery abundance (Figure 6A). However, inhibition of CDK8 with BI-1347 (referred to as CDK8i)^95^ led to significant changes in DDR: ATM, DNA-PKcs and Chk2 phosphorylation levels were elevated whereas Chk1 phosphorylation levels were reduced (Figure 6A), suggesting that DDR is elevated in the absence of functional CKM. A CDK8 knockout HeLa line (Figure S6A) treated with etoposide to introduce DSBs^96^ also caused elevated ATM and Chk2 phosphorylation (Figure S6B), indicating that CKM’s role in regulating DDR is conserved from yeast to mammals.

**Figure 6.**
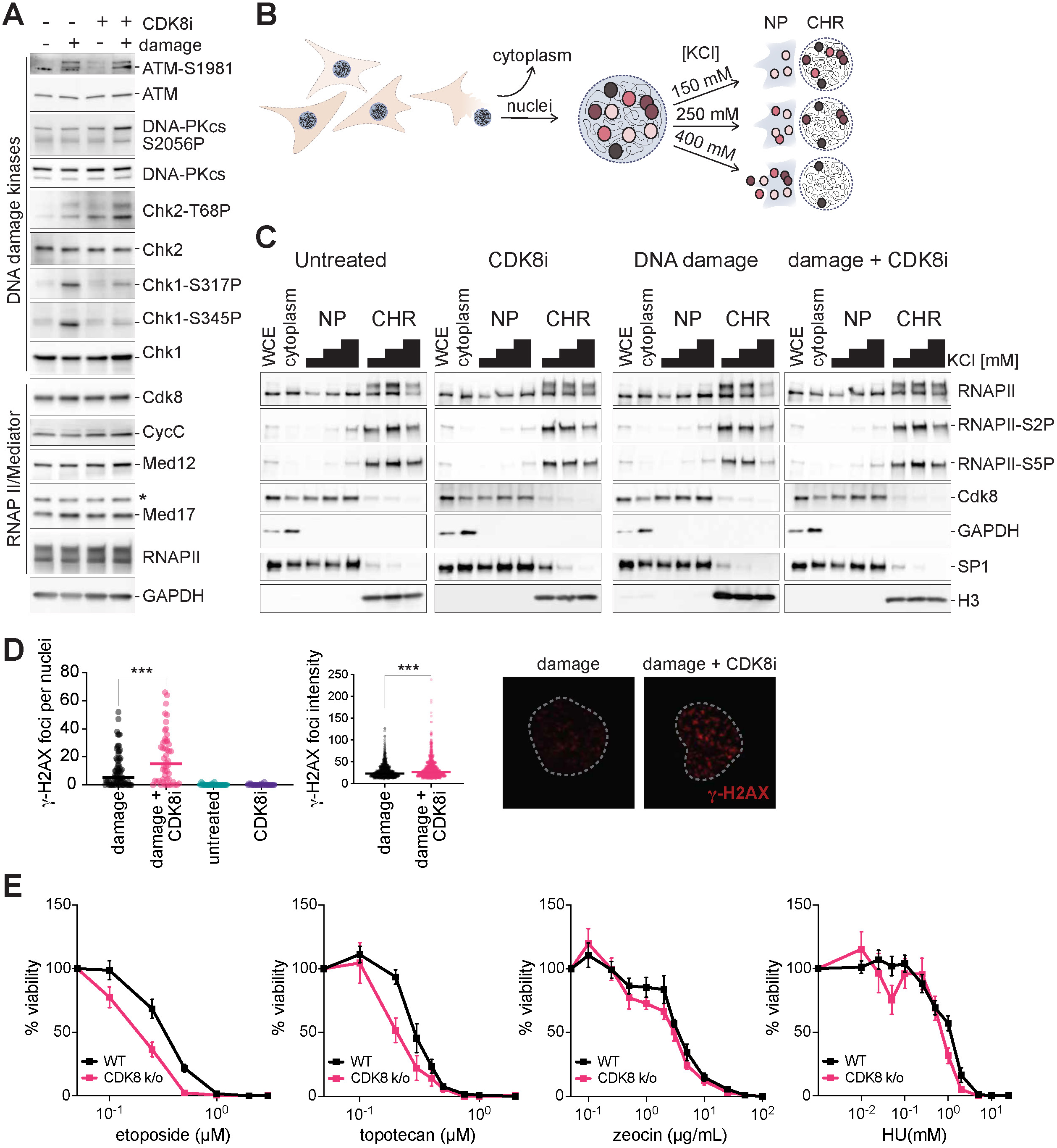
CKM inhibition impedes DNA damage signaling in mammalian cells. **A.** DDR in DivA cells, 4 hours after 4-hydroxytamoxifen (4-OHT) treatment to induce DSBs, with and without CDK8 inhibitor BI-1347, assayed by Western blotting for various DDR markers. **B.** Nuclear fractionation workflow. **C.** Nuclear fractionation of DiVA cells 4 hours after 4-OHT treatment. WCE: whole cell extract, NP: nucleoplasm, CHR: chromatin. GAPDH: cytoplasmic marker, Histone H3: chromatin marker and SP1: nucleoplasmic marker. **D.** γ-H2AX foci number (left) and foci intensity (right) in U2OS DIvA cells with or without CDK8 inhibitor, 4 hours after 4-OHT treatment, immunostained with γ-H2AX (representative images on the right, lines represent median). **D.** Sensitivity of CDK8 knockout HeLa cells to etoposide, TPT, zeocin and HU by clonogenic survival assay (n> 4; error bars represent S.E.M.).

Our findings in yeast show that CKM modulates DDR by reducing active transcription. To test whether CKM has an analogous role in metazoans, we assayed changes in RNAPII chromatin enrichment after DSB induction by salt-fractionation of cultured cell nuclei. Tightly chromatin-bound proteins can only be solubilized with high-salt buffers, whereas proteins that are weakly bound can be readily extracted to nucleoplasm with isotonic buffers (Figure 6B). Western blot analysis of salt-fractionated U2OS DIvA nuclei show that following damage, actively transcribing RNAPII marked by S2 and S5 phosphorylation are more readily extracted from chromatin at high salt concentrations, compared to the untreated cells (Figure 6C). These findings suggest that DSBs also antagonize active transcription in mammalian cells, as observed in yeast. Upon CKM inhibition, a significant portion of actively transcribing RNAPII pools were still retained in high salt chromatin fractions, indicating increased transcriptional activity. We observe the same trend in HeLa cells following siRNA knockdown of CDK8 (Figure S6C, S6D), indicating that CKM limits the chromatin retention and/or recruitment of RNAPII following damage in mammals, akin to our observations in yeast.

In yeast, CKM depletion alters DNA damage foci, as measured by Ddc2 fluorescence (Figure 5C), ChIP analysis of the transcriptional machinery (Figure 5A) and by γ-H2AX spreading at DSBs (Figure 5B). Our analysis of the γ-H2AX foci in U2OS DIvA cells mirror these conclusions: both the γ-H2AX foci number and the γ-H2AX focus intensity are elevated in CKM-inhibited cells (Figure 6D), highlighting CKM’s conserved role in modulating DSB sites.

In yeast, CKM mutants are particularly sensitive to DSB-inducing reagents (Figure 2I). To investigate the effect of CKM loss on DNA damage sensitivity in mammalian cells, we performed a clonogenic survival assay with CDK8 knockout HeLa cells along with a clonogenically-selected WT control. CKM knockout cells conferred increased sensitivity to etoposide and topotecan, which induce DSBs via topoisomerase poisoning. However, CKM knockout lines did not confer statistically significant sensitivity to zeocin, which generates both single-strand and DSBs, or HU which stalls replication forks and exposes single strand DNA. Taken together, these findings that, as in yeast cells, CKM regulates DDR specifically upon DSBs.

## Discussion

Here, we identified a previously uncharacterized role for the CKM of Mediator in transcriptional attenuation in response to DSBs. In the absence of CKM, global transcription is modestly, and cells are defective in mitotic re-entry despite being proficient in DNA repair. An RNAPII mutant with elevated transcriptional output recapitulates the mitotic re-entry defect of CKM mutants, whereas slow RNAPII mutants can rescue CKM loss of function, indicating that transcriptional attenuation by CKM is necessary for mitotic re-entry. Upon DNA damage, CKM interferes with core Mediator-RNAPII binding in a kinase activity dependent manner, effectively limiting transcription initiation. In addition, CKM antagonizes the recruitment of RNAPII to break sites to impede pervasive transcription which could lead to mutagenic DNA repair events. Importantly, we find that CKM’s role in regulating DDR is conserved from yeast to humans. In both contexts, CKM inhibition leads to exaggerated DDR, sensitivity to DSB-inducing reagents and increased γ-H2AX at DSBS. Furthermore, CKM inhibition or deletion also fail to limit the chromatin localization of RNAPII in response to damage in human cell lines. Collectively, our data argue that CKM is a critical regulator of DDR is conserved from yeast to mammals.

We find that a single irreparable DSB, which triggers an extended G_2_/M arrest, attenuates global transcription in budding yeast through a distinct mechanism that previously studied pathways. HU-induced replication stress has also been shown to reduce transcription^27,29^. As previously reported for HU^29^, we show that synthetic upregulation of transcription following a single DSB also further sensitizes cells to damage. However, the mechanistic details of the transcriptional repression in response to DSBs and HU are fundamentally different. First, unlike a DSB which is confined to a single genomic locus, HU-induced replication stress directly interferes with transcription by causing widespread transcription-replication conflicts. Second, HU-induced stress triggers proteasomal depletion of RNAPII. Albeit through a distinct pathway, UV damage has also been shown to downregulate transcription^97^ through depletion of RNAPII^30^. In contrast to both of these mechanisms, a DSB does not alter the abundance of the core transcriptional machinery. Rather, CKM interferes with transcription initiation in response to a DSB by interfering with core Mediator-RNAPII binding. Furthermore, the transcriptomic changes to HU, MMS and an enzymatically induced DSB are considerably different. These disparities may result from the differences in the cell cycle arrest induced by these reagents as well as the specific DDR kinases enforcing the arrest. While DSBs antagonize transcription during prolonged G_2_/M arrest, HU induces a transcriptional response predominantly in S phase^27,29^. Therefore, various types of DNA damage inhibit transcription through different pathways with and CKM playing a distinct role in transcriptional downregulation upon DSBs.

Cdk8, the kinase subunit of CKM, belongs to the cyclin-dependent kinase family of proteins which regulate cell cycle^98^. Canonical cyclin-dependent kinases are regulated throughout the cell cycle by the abundance of their respective cyclins^98^, yet CycC, the cyclin of Cdk8, is non-cycling^98^. Although Cdk8 modulates G_2_/M transition in yeast^99^ and in mammals^100,101^, implying that Cdk8 is regulated by cell cycle, the underlying molecular mechanisms are poorly understood. Here, we present evidence to suggest that both CKM abundance and kinase activity are elevated soon after G_2_/M arrest and are diminished as cells progress into mitosis. Based on these findings, we posit that cell cycle or DDR-specific kinases could differentially modulate CKM to stimulate its activity in a cell cycle-specific manner. Supporting this, CKM, especially the intrinsically disordered region (IDR) of Med13, is extensively phosphorylated in response to DNA damage^102,103^. Strikingly, the Med13 IDR is responsible for occluding RNAPII binding to core Mediator^38^. The IDR phosphoacceptor sites could also be targeted by DDR kinases or by CKM itself to fine-tune CKM-core Mediator binding, and therefore transcription. Alternatively, DDR-dependent phosphorylation of CKM subunits could target them for degradation, as shown for Med13^104^ and for CycC^105^ under oxidative stress.

CKM and RNAPII compete for the same binding face on the core Mediator, therefore CKM is thought to occlude RNAPII binding via steric hindrance^38,42–45^. In addition, recent *in vitro* work shows that CKM kinase activity facilitates the dissociation of CKM-core Mediator complex^42^, likely through the phosphorylation of this shared binding interface. Consistent with this kinase-dependent dissociation mechanism, we find that stimulation of CKM kinase activity by damage reduces core Mediator-RNAPII binding, while CKM inhibition enhances this interaction. Upon damage, CKM abundance is reduced as a function of time; yet, this reduction does not lead to a corresponding increase in core Mediator-RNAPII binding, as would be expected with a steric hindrance model. Based on these findings, we postulate that the binding interface of core Mediator, which is shared by CKM and RNAPII, could be post-translationally altered in response to DNA damage by CKM, rendering it refractory to binding and effectively inhibiting core Mediator-RNAPII complex formation. In agreement with this model, CKM inhibition following DNA damage enhances core Mediator-RNAPII binding. We also note that this CKM kinase-dependent dissociation could enable rapid recovery from the damage-dependent transcriptional arrest simply by dephosphorylation of core Mediator.

We find that CKM’s mitotic re-entry defect is rescued by downregulation of transcription, either by depletion of preinitiation complex or reducing RNAPII polymerization rate. Additionally, a hyperactive RNAPII mutant is also defective in mitotic re-entry, indicating that elevated transcription is detrimental for DDR. As CKM depletion does not markedly alter specific cell cycle or DDR transcripts which could account for the mitotic re-entry defect of CKM mutants, we postulate that the detrimental effect of transcription on DDR is indirect. For instance, increased transcription could expose single-stranded DNA, which could then serve as a template for the recruitment and activation of DDR kinases. In agreement with this, we detect increased γ-H2AX spreading within gene bodies upon CKM inhibition while the magnitude of γ-H2AX signal around DSBs remains largely unaltered. These findings suggest that in response to DNA damage, CKM might function to reduce the amount of exposed single-strand DNA which might trigger spurious activation of DDR kinases.

Our tetrad analysis with *cdk8-kd*, *rad53-kd* and *mec1-kd* strains as well as the E-MAP data point to a possible DNA damage and Mec1-independent interaction between *RAD53* and CKM under basal conditions, which can partially be attributed to elevated histone expression. However, the molecular mechanisms underlying the genetic interaction between Rad53-CKM under basal conditions remains to be investigated. Rad53 binds to nearly 20% of yeast promoters and contributes to their transcriptional regulation under basal conditions^18^. CKM could be involved in the transcriptional regulation of these Rad53-bound genes, which in turn could affect growth. Future work will focus on the interaction between these two kinases.

CKM structure and transcriptional function is largely conserved from yeast to mammals^35–38^. Here, we find that CKM’s role in DDR regulation is also conserved from yeast to humans. As in yeast, CKM inhibition or depletion upregulates DDR in human cells. Akin to our findings in yeast, CKM mediates RNAPII chromatin eviction in response to DNA damage and provides resistance to double-strand break inducing reagents. Consonant with our ChIP-seq analysis in yeast, we find that CKM inhibition in mammalian cells leads to increased γ-H2AX foci. Intriguingly, CKM was identified as an interaction partner of with DNA-dependent serine threonine kinase DNA-PKcs, which is a regulator of mammalian DDR^41^. Moreover, CKM is implicated in various cancers^95,101,106–108^; however, whether CKM’s role in malignant transformation is related to its function as a regulator of DDR remains to be resolved. Taken together, these observations suggest that CKM acts as a conserved regulator of transcriptional repression upon DSBs and presents a critical node in DDR regulation across eukaryotes.

### Resource availability Lead contact

Requests for additional information, resources or raw data should be directed to and will be fulfilled by the Lead Contact.

## Materials availability

Cell lines, yeast strains and plasmids generated in this study are available upon request.

## Data and code availability

RNA-seq, PRO-seq and ChIP-seq data generated for this study will be made available at the Gene Expression Omnibus (GEO) after peer review (GSE296170, GSE296218, GSE296384). Remainder of the raw data reported in this paper are available upon request. No original code was generated in this study. The code and software used to analyze data are listed in STAR Methods.

## Acknowledgements

This work was supported by the NIH-NRSA (NIGMS, F32GM145156) and the University of Chicago Women’s Board Grant Fund to G.M.; NIH (NIGMS, R35GM145373) and the University of Chicago ITM Subsidy Award to A.J.R.; NIH (NIGMS, R35GM127029) to J.E.H. and NIH (NIGMS, R01GM084279 and NCI U54 CA274502) to N.J.K. We thank A. Mohan and Q. Wu from Haber Lab (Brandeis) for generating yeast strains; J. Grubb and D. Bishop from Bishop Lab; B. Budke from Connell Lab for suggestions, reagents and advice; S. Mukherjee, H.C. Lee and D. Pincus’ lab at the University of Chicago for sharing equipment. We thank the University of Chicago Genomics (RRID: SCR_019196), Integrated Light Microscopy (RRID: SCR_019197), Cytometry and Antibody Technology (RRID: SCR_017760) and Sanger Sequencing Cores (RRID: SCR_025314). We thank A. Kainth for his feedback on the manuscript. The HeLa cell line was established from the tumor cells of Mrs. Henrietta Lacks in 1951 without her consent and has been instrumental for scientific research. We are grateful to Mrs. Henrietta Lacks and her surviving family for their contributions.

## Author contributions

Conceptualization and methodology, G.M., N.J.K., J.E.H. and A.J.R.; validation, analysis, investigation, visualization, G.M; data curation, G.M, S.B.; Writing-original draft, G.M.; Writing-review & editing, G.M., J.E.H., A.J.R; Funding acquisition, G.M., N.J.K., J.E.H., A.J.R.; Supervision, N.J.K., J.E.H., A.J.R.

## Declaration of interests

The Krogan Laboratory has received research support from Vir Biotechnology, F. Hoffmann-La Roche, and Rezo Therapeutics. N.J.K. has a financially compensated consulting agreement with Maze Therapeutics. N.J.K. is the President and is on the Board of Directors of Rezo Therapeutics, and a shareholder in Tenaya Therapeutics, Maze Therapeutics, Rezo Therapeutics, and GEn1E Lifesciences.

## Supplemental Excel File Titles

**Supplemental Table 1.** CC scores obtained from E-MAP screen

**Supplemental Table 2.** Plasmids used in this study

**Supplemental Table 3.** Yeast strains used in this study

**Supplemental Table 4.** ssDNA repair templates used for CRISPR/Cas9-targeting

**Supplemental Table 5.** Antibodies used in this study

**Supplemental Table 6. q**PCR primers used in this study

## Supplemental Figure Titles and Legends

**Supplementary Figure 1, related to Figure 1.**
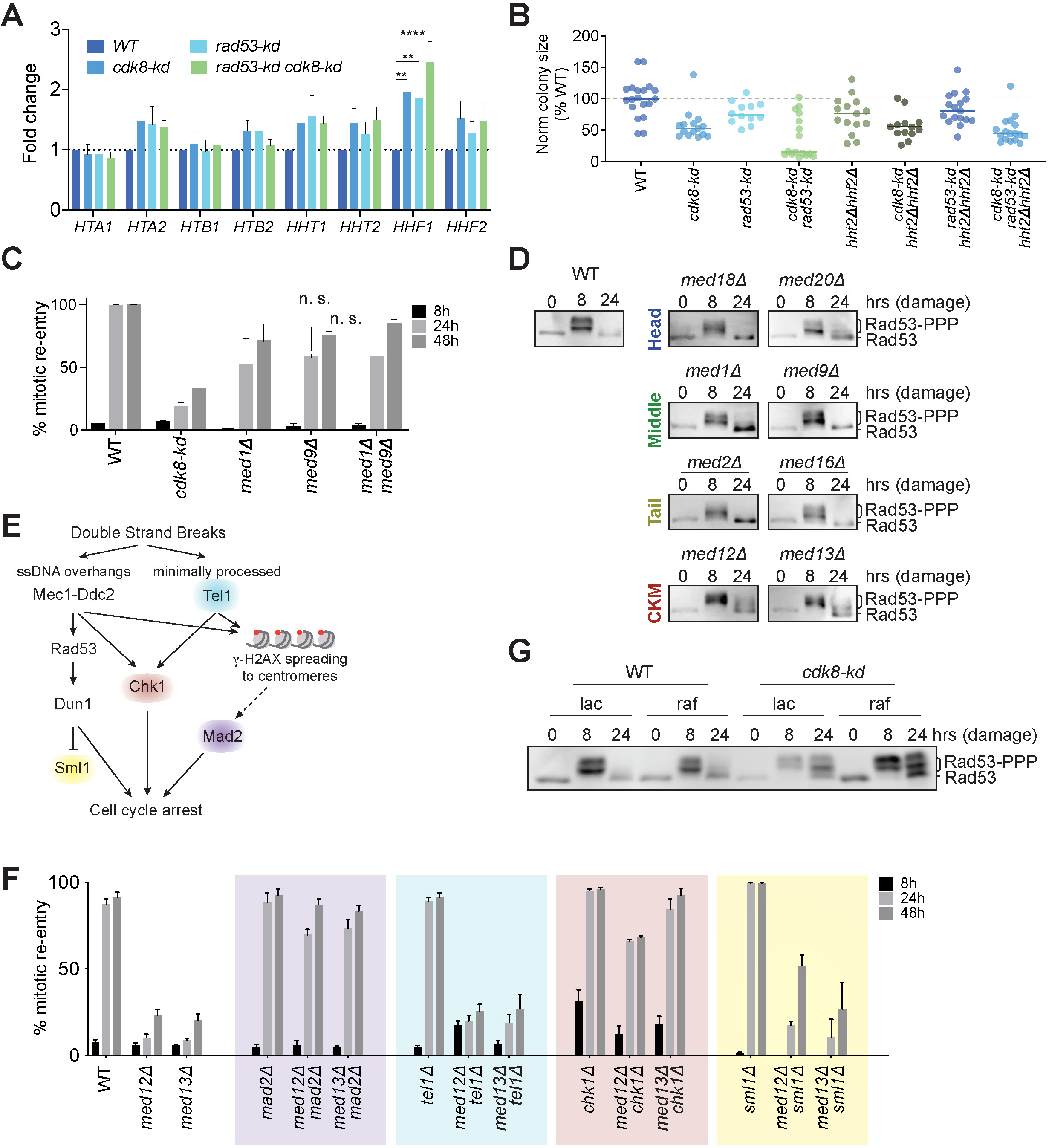
CKM acts on the Mec1 branch of the DDR to promote mitotic re-entry. **A.** Relative abundance of individual histone gene transcripts measured by RT-qPCR in WT, *cdk8-kd*, *rad53-kd sml1*1 and *cdk8-kd rad53-kd sml1*1 strains (n=4; error bars represent S.E.M.; \*\*\**p*<0.0001; \*\**p*<0.01; 2-way ANOVA). **B.** Normalized colony sizes from WT control strain crossed to *cdk8-kd rad53-kd hht2*1 *hhf2*1 quadruple mutant. All strains contain a compensatory *sml1*1 mutation to ensure the viability of *rad53-kd*. Nearly one-third of the *cdk8-kd rad53-kd* double mutants dissected from this cross grow to WT levels, possibly due to suppressor mutations, masking the effect of *hht2*1 *hhf2*1. On the other hand, all the *cdk8-kd rad53-kd* double mutants obtained from the cross represented in Figure 1D are severely defective in growth. **C.** Percent mitotic re-entry following a single DNA break in *med1*1, *med9*1 and *med1*1 *med9*1 mutants. **D.** DNA damage response to a single irreparable DNA break as measured by Rad53 hyperphosphorylation in Mediator head, middle, tail and CKM mutants. **E.** A simplified diagram of the DNA damage response (DDR) pathway in budding yeast, adapted from Harrison *et. al*^110^. Minimally processed DNA breaks stimulate Tel1 kinase, whereas single stranded DNA exposed by 5’ to 3’ resection leads to the recruitment of Mec1 kinase to DNA breaks via its binding partner Ddc2. These two upstream kinases activate the downstream kinase Chk1. The activation of downstream effector kinase Rad53 in response to double-strand breaks is dependent on Mec1. Rad53 then stimulates Dun1 kinase, which suppresses Sml1 to upregulate dNTP levels. Mec1 and Tel1 kinases phosphorylate H2A (which is referred to as γ-H2AX). The spreading of γ-H2AX at later stages of DNA damage-induced cell cycle arrest also activates spindle assembly proteins, including Mad2; which contributes to the cell cycle arrest. **F.** Percent mitotic re-entry in CKM (*med12*1 or *med13*1) and DDR (*mad2*1, *tel1*1, *chk1*1 and *sml1*1) double mutants. **G.** DNA damage response of WT and *cdk8-kd* cells grown in YEP-lactate and YEP-raffinose to a single irreparable break as measured by Rad53 hyperphosphorylation.

**Supplementary Figure 2, related to Figure 3.**
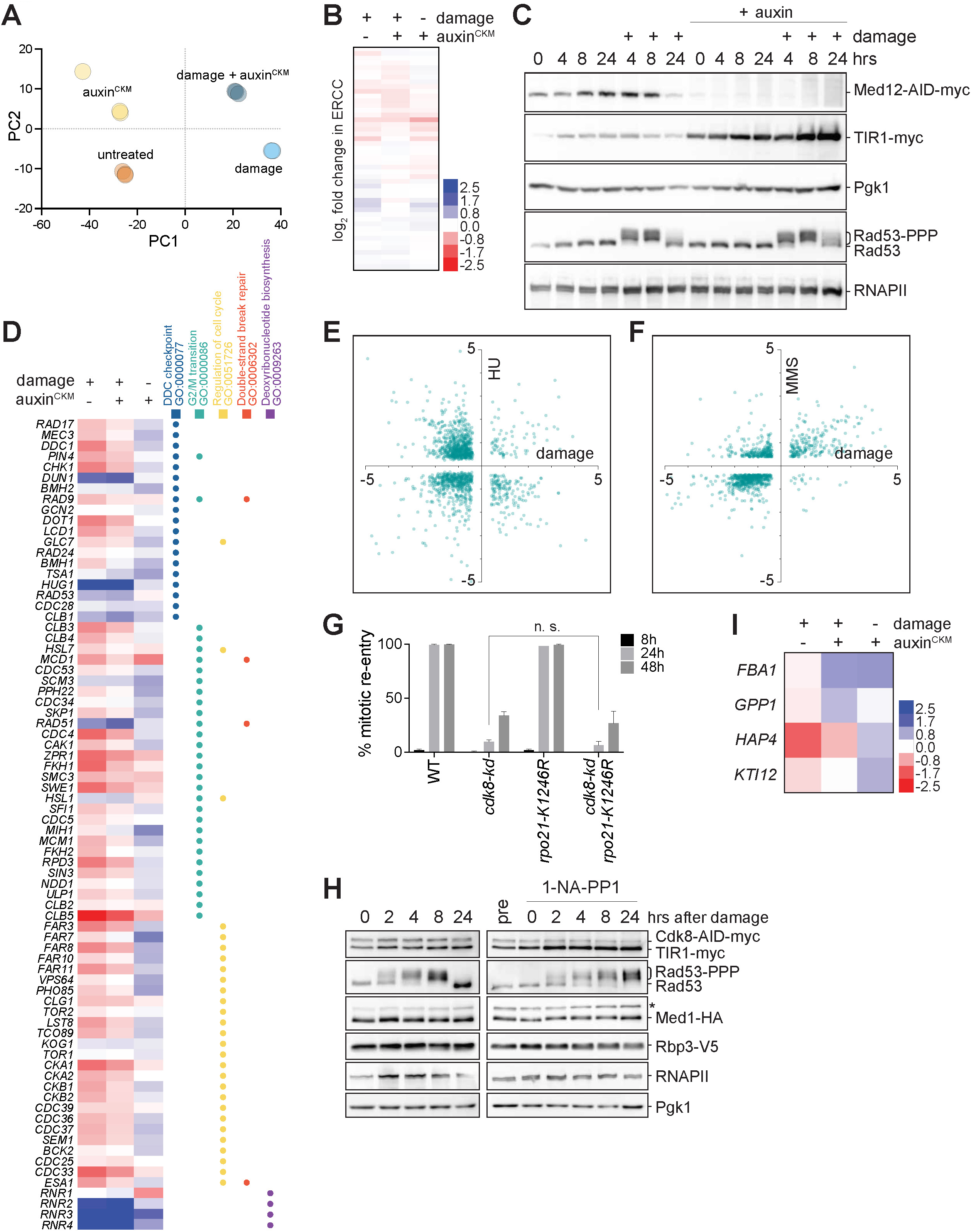
DNA damage and CKM activity do not alter RNAPII protein levels or DNA damage-related transcripts. **A.** Principal component analysis for bulk RNA sequencing analysis shown in Figure 3A (n=3) demonstrates robust replicate measurements. **B.** Log_2_ fold change in external RNA (ERCC) spike-in controls for the bulk RNA sequencing analysis shown in Figure 4A, as a background control for sample preparation, library preparation or sequencing artifacts. The spike-in transcript levels are comparable across samples (n=3), accommodating for global effects. **C.** Cell lysates collected side-by-side with the samples for bulk RNA sequencing (Figure 3A) blotted for myc (to detect Med12-AID-myc and TIR1-myc), Pgk1, RNAPII and Rad53 to assay the DNA damage response. **D.** Log_2_ fold change in expression of genes which are involved in DNA damage response and cell cycle extracted from relevant gene ontology terms^111^, from the differential expression analysis shown in Figure 3A. **E.** Comparison of significant (p_adj_ > 0.05) log_2_ fold change in expression following 45 minutes of HU treatment, adapted from Sheu *et al*^18^ to significant log_2_ fold change in expression 8 hours after the induction of DNA damage. **F.** Comparison of significant log_2_ fold change in expression following 1 hour of 0.03 % methylmethane sulfonate (MMS) treatment, adapted from Jaehnig *et* al^19^ to significant log_2_ fold change in expression 8 hours after the induction of DNA damage. **G.** Percent mitotic re-entry following a single DNA break in *rpo21-K1246R* and *cdk8-kd* mutants after induction of DNA damage. **H.** Cell lysates collected side-by-side with the samples used for chromatin immunoprecipitation experiment shown in Figure 3D, blotted for myc (to detect Cdk8-AID-myc and TIR1-myc), HA (to detect Med1-HA), V5 (to detect Rbp3-V5), RNAPII, Pgk1 and Rad53 to assay the DNA damage response. **I.** Log_2_ fold change in expression of three control loci picked to assay RNAPII enrichment in Figure 3D, extracted from the differential expression analysis shown in Figure 3A.

**Supplementary Figure 3, related to Figure 3.**
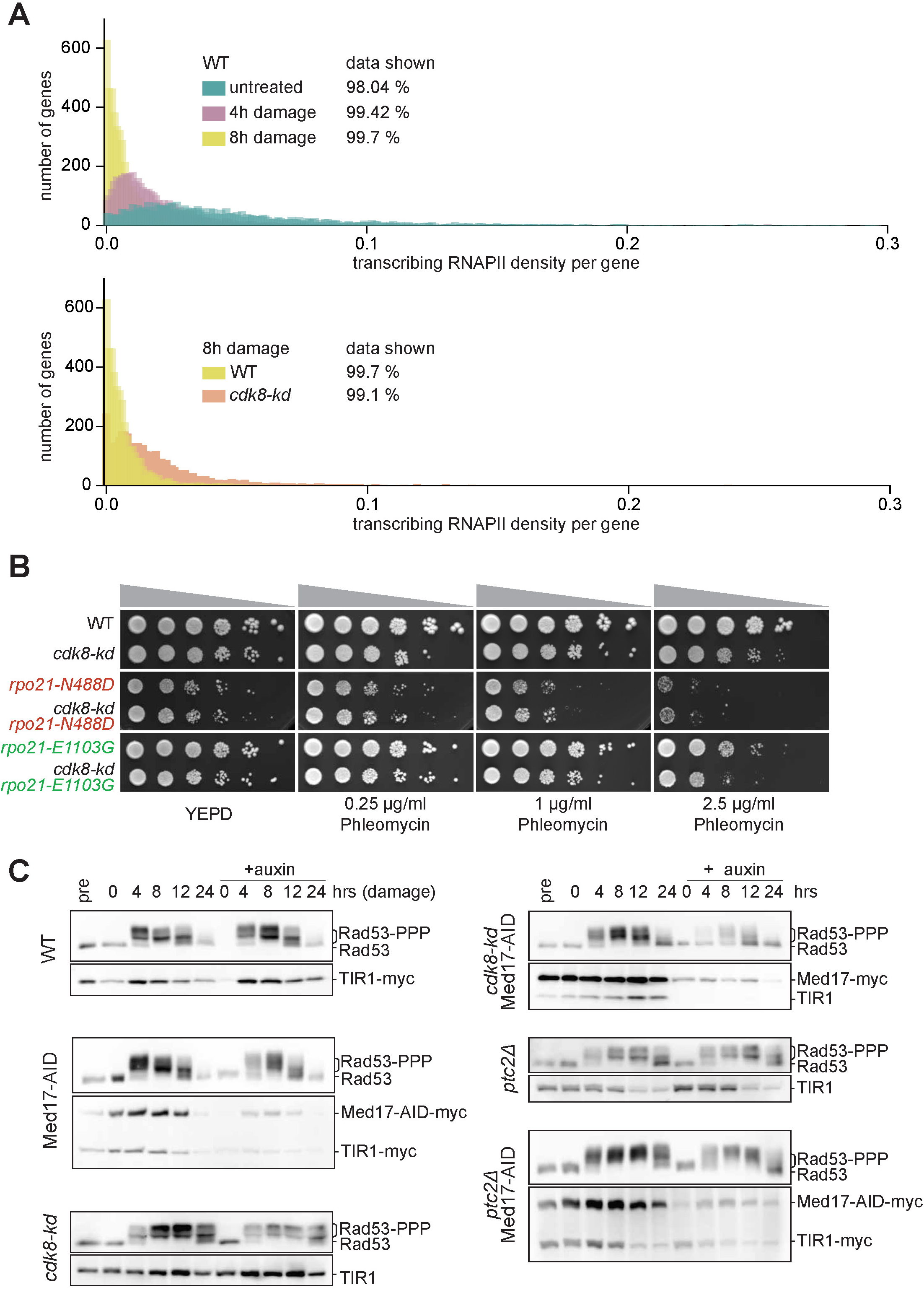
Transcriptional output dictates DNA damage response. **A.** Transcribing RNAPII density per gene as measured by PRO-seq in WT and *cdk8-kd* cells following DNA damage. Truncated versions of these histograms are represented in Figure 3E and 3F. To maintain a scale which still allows the visualization of the data distribution, the genes which have an RNAPII density above 0.3, which are not visible with Y-axis scaling, were discarded from the plot. The percentage of the dataset plotted for this analysis for each strain is indicated on corresponding graphs. **B.** Sensitivity of *rpo21-N488D* (slow; red) and *rpo21-E1103G* (fast; green) RNAPII mutants with or without *cdk8-kd* to various phleomycin concentrations. **C.** DNA damage response to a single irreparable DNA break as measured by Rad53 hyperphosphorylation following the auxin-induced depletion of essential core Mediator subunit Med17 in *cdk8-kd* and *ptc2*1 mutants.

**Supplementary Figure 4, related to Figure 4.**
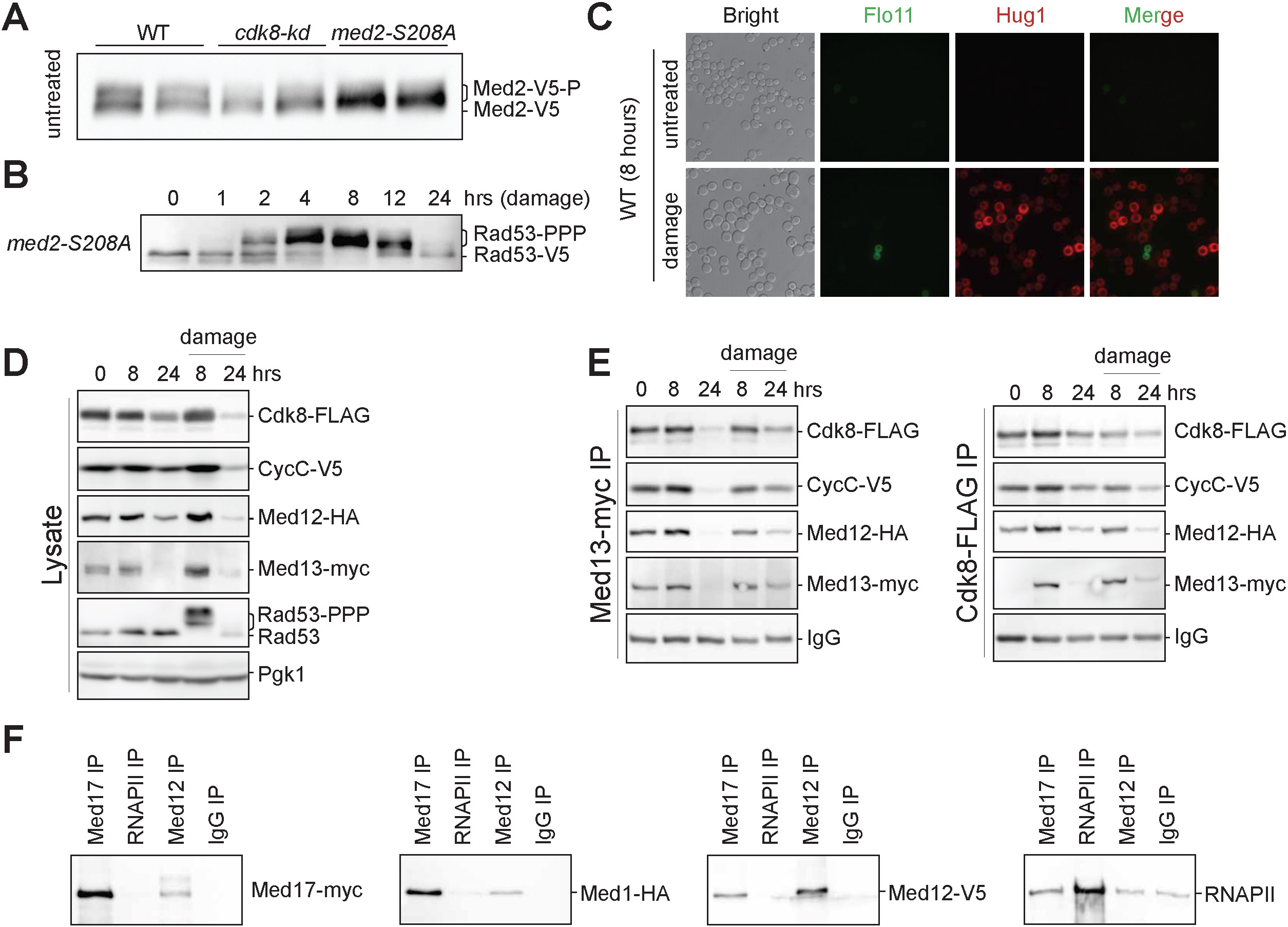
CKM stoichiometry remains the same after DNA damage, while CKM-core Mediator binding is altered in response to damage and CKM inhibition. **A.** Med2 phosphorylation in untreated WT, *cdk8-kd* and *med2-S208A* mutants, as detected by a hypomobility shift on a Western blot. Samples were ran side by side for comparison of the Med2 hypomobility shift in these strains. **B.** DNA damage response in *med2-S208A* mutant as measured by Rad53 hyperphosphorylation following the induction of a single irreparable break. **C.** Flo11::mNeonGreen or Hug1::mScarletI in WT, following 8 hours of DNA damage. **D-E.** CKM stoichiometry changes in WT cells with Cdk8-FLAG, CycC-V5, Med12-HA and Med13-myc epitope tags, assayed by Med13-myc (middle) or Cdk8-FLAG (right) immunoprecipitation. Lysate samples (**D**) were blotted for Rad53 to infer the DNA damage response and for Pgk1 as loading control. **F.** Med17, RNAPII and Med12 interactions assayed by immunoprecipitation in the presence of benzonase to degrade DNA and RNA, followed by Western blotting. IgG was used as a negative control to determine nonspecific binding to beads. These assays were carried out side by side with the immunoprecipitation experiments shown in Figure 4F and Figure S4F.

**Supplementary Figure 5, related to Figure 5.**
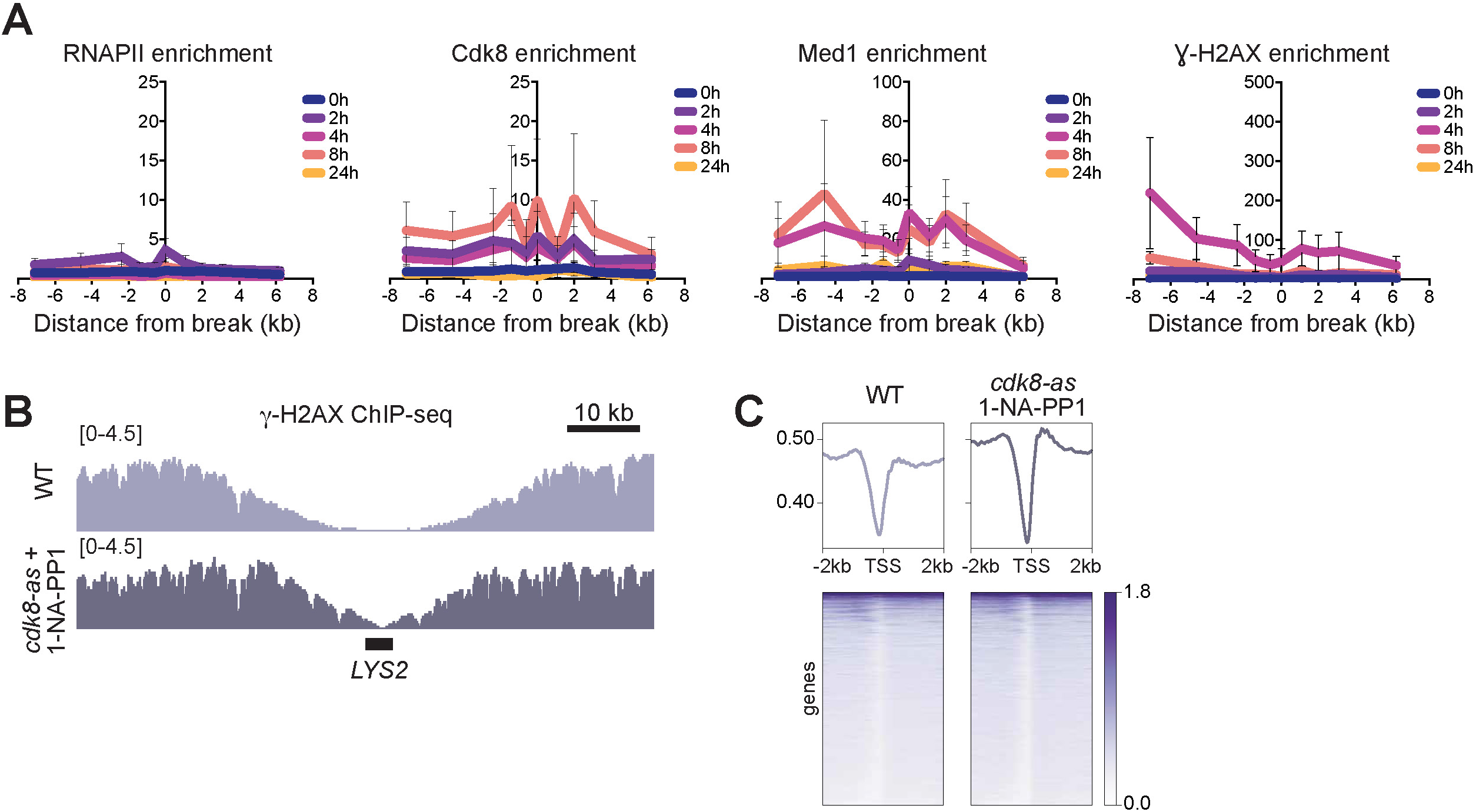
CKM localizes to DNA breaks together with core Mediator. **A.** The WT strain used for the ChIP-qPCR experiment shown in Figure 5A has another DNA break site engineered at the *LYS2* locus. Fold enrichment of RNAPII (Rbp3-V5), Cdk8 (Cdk8-myc), core Mediator subunit Med1 (Med1-HA) and γ-H2AX around *LYS2* locus as a function of time after DNA damage, measured by chromatin immunoprecipitation recapitulates nearly the same results observed for the DNA break at *MAT* locus shown in Figure 5A. **B.** γ-H2AX around the DNA break at *LYS2* locus 8 hours after the induction of DNA damage as measured by ChIP-seq. **C.** Aggregate plot of normalized γ-H2AX ChIP signal within open reading frames, 8 hours after the induction of DNA damage. TSS indicates transcription start sites and TES indicates transcription end sites.

**Supplementary Figure 6. related to Figure 6.**
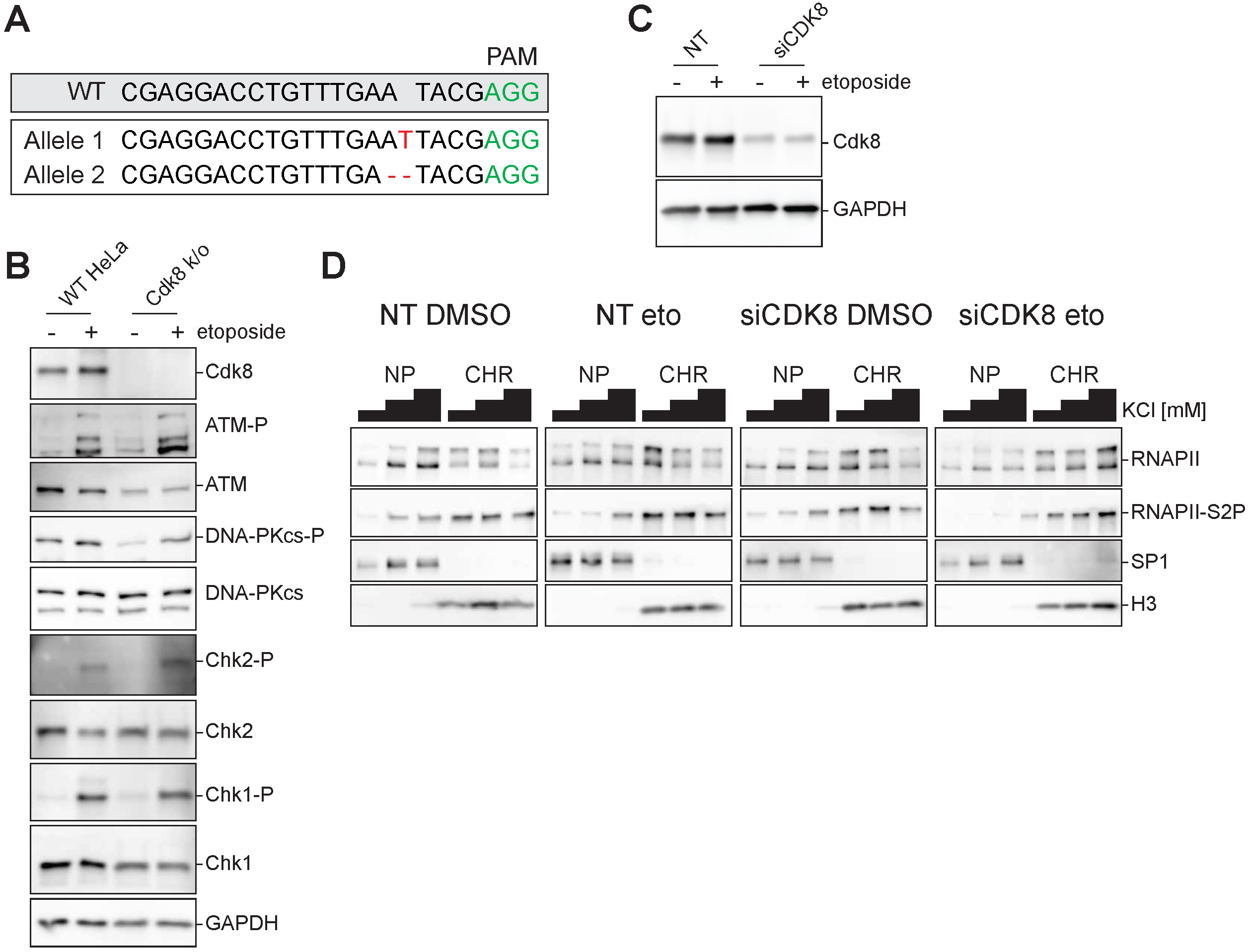
CKM modulates chromatin recruitment of RNAPII in response to DNA damage in mammalian cells. **A.** Genotype of CDK8 HeLa knockout line. WT sequence is indicated above, and the PAM sequence used for the CDK8 gRNA design is indicated in green. The mutations detected in CDK8 ORF, illustrated in red, both lead to out-of-frame frameshifts. **B.** The DNA damage response in CDK8 knockout HeLa cells assayed by phospho-specific antibodies against DNA damage-dependent phosphorylation of key DDR kinases. Samples were also blotted with a CDK8-specific antibody to confirm the CDK8 knockout in this cell line. **C.** Confirmation of CDK8 knockdown with siRNA in HeLa cells by Western blotting. NT: nontargeting control siRNA. **D.** Nuclear fractionation of HeLa cells (with or without siCDK8) 6 hours after etoposide treatment to measure the chromatin occupancy of active transcriptional complexes (RNAPII). Bars above the blots represent the increasing salt concentration in fractionation buffers (150 mM, 250 mM, 400 mM). NP: nucleoplasm, CHR: chromatin.

## STAR Methods

### Experimental Model and Subject Details

Yeast strains were freshly thawed from frozen stocks and grown at 30^°^C using standard practices. Plasmids and yeast strains used in this study are listed in Table S2 and S3 respectively. All yeast strains used in this study are derivatives of S288c. ORF deletions are introduced either through one step PCR homology cassette amplification^112^ or by using CRISPR/Cas9 as previously described^113^. The sequences of template DNAs used for gene targeting experiments are listed in Table S4. Epitope-tagged strains were created either by using PCR-amplified homology cassettes^114^ or by using CRISPR/Cas9 gene targeting methods^113^. The AID-tagged strains were constructed as previously described^115^. HeLa cells were obtained from ATCC. DIvA cells^88^ (AsiSI-ER-U2OS) were a kind gift from Dr. Gaëlle Legube (Centre de Biologie Intégrative, Toulouse, France). All mammalian cell lines were maintained under sterile conditions and tested regularly for contamination by DAPI-staining combined with microscopy and with PCR-based Venor GeM Mycoplasma Detection Kit (Sigma-Aldrich, catalog number # MP0025). The mycoplasma test results are available upon request.

### Yeast Media and Treatments

Yeast strains were grown in YEP-dextrose (1% yeast extract (w/v), 2% peptone (w/v), 2% dextrose (w/v), 0.004% adenine (w/v), pH5.5). For experiments involving a galactose induction, cells were inoculated in YEP containing 3% lactic acid (v/v, Fisher Scientific, cat # A162-1) or 2% raffinose (w/v, Sigma-Aldrich, cat # 83400) and treated with a final concentration of 2% galactose (w/v, Sigma-Aldrich, cat # G0750). For the inducible degradation of the proteins, the cultures were treated with 100 μg/mL indole-3-acetic acid (IAA) (Sigma-Aldrich, cat # I2886). For inhibition of Cdk8 kinase activity in *cdk8-as* (*cdk8-Y236G*) mutant, cells were treated with a final concentration of 20 μM 1-NA-PP1 (Cayman Chemical, cat # 10954-10) unless otherwise indicated. To induce a G_2_/M arrest by microtubule inhibition, cells were treated with 10 μg/mL of nocodazole (US Biological, cat # N3000). Temperature-sensitive conditional mutant *cdc13-1* was grown at permissive temperature (25^°^C) and shifted to nonpermissive temperature (33^°^C) to induce DNA damage.

### Epistatic Miniarray Profiling Screen (E-MAP)

The E-MAP screen^33^ was performed as explained as before^56^. The yeast strains used for the E-MAP assay are listed in Table S3. The E-MAP scores obtained the mutants were first clustered by using Cluster^116^ and then plotted by TreeView^117^. The CC scores from the E-MAP experiment are listed in Table S1.

### Tetrad Assays

Strains containing *mec1-kd sml1*1 or *rad53-kd sml1*1 mutations were crossed to *cdk8-kd sml1*1 strains. Resulting zygotes were picked by micromanipulation and grown at 30°C on YEP-dextrose plates. Grown diploids were sporulated on SPM plates (3 % potassium acetate (w/v), 0.02 % raffinose (w/v), 2 % agar (w/v) and 0.012 % of essential amino acid powder (w/v)) overnight. The next day, cells grown on sporulation plates were resuspend in 20 μL of filter-sterilized ZK buffer (25 mM Tris-Cl pH 7.5; 0.8 M KCl) and treated with zymolyase 100T (US Biological, cat # Z1004) at final concentration of 0.4 mg/mL. The cells were incubated at 37^°^C for 5 minutes in zymolyase to digest the cell wall and then spread on a YEP-dextrose plate for dissection. Dissected tetrads were grown at 30^°^C until colonies form. Once grown, tetrad colonies were genotyped via PCR, sequencing and antibiotic resistance. Tetrad colony plates were imaged by ImageQuant 800 (Amersham). The colony sizes of tetrads were quantified by using FiJi^118^, normalized to WT and plotted by using Prism 10 (Graphpad).

### Yeast RNA extraction, cDNA preparation and RT-qPCR

10 mL of cells were grown to early exponential phase in YEP-dextrose were harvested by centrifugation at 4^°^C. The cell pellets were washed twice with ice cold water and stored at -80^°^C. RNA from cell pellets were extracted using YeaStar RNA kit (Zymo Research, cat # 11-329), treated with Turbo DNase (Invitrogen, cat # AM2238) and diluted to a final concentration of 100 - 150 ng/μL. 10 μL of RNA was used for cDNA prep by using MMLV reverse-transcriptase (Lucigen, cat # RT80125K), and the reactions were supplemented with RiboGuard RNase inhibitor (Biosearch Technologies, cat # RG90925) to prevent RNA degradation. Resulting cDNA was used to set up qPCRs with primers specific to each histone gene as detailed below. The primer sequences are listed in Table S6.

### Real time quantitative PCR

3 identical reactions were set up for each DNA and primer pair with 2X PowerUp SYBR Green Master Mix (Applied Biosystems, cat # A25741) using BioRad CFX384 thermocycler. The Cq values for at least 3 technical replicates were averaged for each reaction, normalized to a control locus, the input, and the untreated sample for ChIP experiments. For measuring changes in transcript abundance, Cq values were normalized to a control locus and then to the WT control. Results from at least 3 independent experiments were plotted by using Prism 10 (Graphpad). The primer sets used for the qPCR assays are listed in Table S6.

### Mitotic re-entry assays with micromanipulation

Liquid cultures were grown overnight in YEP-lactate. 50 individual G_1_ cells were micromanipulated on YEP-galactose plates. At various time points, number of cells were scored as a ratio of cells that are past the G_2_/M arrest stage and rebudded to total number of live cells, as previously described^3^. For assays with *cdc13-1* strains, cells were micromanipulated on YEP-dextrose plates and grown at permissive temperature (25^°^C) as control, or nonpermissive temperature (33^°^C) to induce DNA damage. For each mitotic re-entry assay, results from at least 3 independent experiments were plotted by using Prism 10 (Graphpad). All mitotic re-entry assay error bars represent S. E. M., *** *p*<0.001, tested by 2-way ANOVA.

### HO-induction time course

Overnight YEP-dextrose liquid cultures were washed 3 times with YEP-lactate and were grown in YEP-lactate for 6-8 hours (nearly two doubling times). This culture was then used to inoculate a larger YEP-lactate culture. When the density of the culture reached early exponential phase (1-10 x 10^6^ cells/ml), DNA damage was induced by addition of galactose to the media at a final concentration of 2% (w/v). For western blot analyses from cell lysates, 50 mL of culture was harvested by centrifugation, washed with 1 mL of 20 % trichloroacetic acid (TCA) (w/v, Sigma-Aldrich, cat # T9159) and snap-frozen on dry ice, stored at -80^°^C. For coimmunoprecipitation experiments, ∼ 500 mg of wet yeast pellet was collected, washed with 1 mL of ice-cold Tris-EDTA buffer (50 mM of Tris pH 8.0, 5 mM of EDTA, pH 8.0) and snap-frozen on dry ice and stored at -80^°^C. For chromatin immunoprecipitation experiments, 45 mL of culture was crosslinked with 1% methanol-stabilized formaldehyde (Sigma-Aldrich, cat # 252549) for 10 minutes at room temperature with gentle shaking. The crosslinking reaction was quenched by addition of glycine at a final concentration of 125 mM and incubation on a platform rocker for 5 minutes at room temperature. Crosslinked cultures were spun down at 4^°^C. Harvested cell pellets were washed 3 times with 1 X TBS (pH 7.4), flash-frozen on dry ice and saved at -80^°^C. For RNA sequencing experiments, culture density was monitored throughout the time course to maintain cultures at early exponential phase. 50 mL of culture was harvested with centrifugation. The cell pellet was washed with 1 mL of ice-cold water, flash frozen in liquid nitrogen and stored at -80^°^C.

### Yeast Sample preparation for Western Blotting

Yeast whole cell lysate samples were prepared as described^119^. Briefly, frozen cell pellets were resuspended in 200 μl of 20 % TCA on ice and lysed with vortexing for 3 minutes with acid-wash glass beads (Sigma-Aldrich, cat # G8772). The glass beads were washed with 200 μl of 5% TCA twice and the crude extract was collected to a new tube. The crude extract was spun down at 3000 rpm for 10 minutes. The TCA supernatant was discarded, and the protein pellet was resuspended in 200 μL of 6X Laemmli buffer (60 mM Tris-HCl, pH 6.8, 2% SDS (w/v), 10% glycerol (v/v), 0.2 % bromophenol blue (w/v), 0.9 % 2-mercaptoethanol (v/v)) and neutralized with 200 μL of Tris (1M, pH 8.0). Prior to loading, the samples were boiled at 95^°^C for 5 minutes and spun down for 2 minutes at maximum speed at room temperature.

### Western blotting

Polyacrylamide gels were run by using BioRad Mini-PROTEAN tetra cell system in 1X SDS-PAGE buffer, made by diluting the 10 X run buffer stock (250 mM Tris, 1.92 M glycine, 1 % SDS (w/v) in 1000 mL of distilled water). Once the desired protein separation was achieved, the proteins were transferred to Millipore Immobilon-PSQ PVDF membrane (cat # ISEQ00010) at 4^°^C by using BioRad Mini Trans-Blot Cell in 1 X transfer buffer containing 100 mL methanol (from 10 X transfer buffer: 250 mM Tris base, 1.92 M Glycine) to a final volume of 1000 mL with distilled deionized water. After transfer, the membranes were blocked in 1 X TBS (for 10 X TBS, 200 mM Tris, 1.5 M NaCl in 1000 mL distilled water, pH 7.4 with concentrated HCl) containing 1 mL of Tween-20 (Fisher Scientific, cat # BP337-500) and 5 % non-fat dry milk (w/v). For phospho-specific antibodies and histone antibodies, 2 % of ECL Blocking Reagent (Amersham, cat # RPN2125, w/v) was used instead of dry milk. After blocking, membranes were blotted with primary antibodies (1:500 - 1:5000 in blocking solution). Membranes were washed 3 times with 1 X TBS-T and then incubated in secondary antibodies (1:5000 - 1:20,000 in blocking buffer). Membranes were washed again with 1 X TBS-T, developed by using Amersham Select Western Blot Detection Reagent (Amersham, cat # RPN2235) and visualized by using GE ImageQuant LAS 4000 Biomolecular Imager. BioRad ImageLab software was used to process the images. Antibodies used for western blotting are listed in Table S5.

### Viability Assays

For gene conversion and single strand annealing viability assays, to avoid the flocculation phenotype of the CKM mutants convoluting the viability results, we employed a micromanipulation-based viability assay. For this, the cells were grown in liquid YEP-lactate overnight and micromanipulated on YEP-galactose plates as described above for the adaptation assays. Viability was calculated as a ratio of microcolonies formed to the total number of cells divided at 24 hours from at least 3 independent experiments. Results were averaged and plotted by using GraphPad (n>3, error bars represent S.E.M.; one-way ANOVA, n.s.). For nonhomologous end joining (NHEJ) viability assays, cells were grown in YEP-lactate overnight. The next day, cultures were gently sonicated to break up flocculent clumps on Branson 2800 Ultrasonic Water Bath (two minutes on two minutes off for 5 cycles), counted and plated either on galactose or dextrose containing plates. The plates were incubated at 30^°^C until visible colonies emerged. NHEJ viability was calculated as the ratio of colonies on galactose plates to colonies on dextrose plates from at least 3 independent experiments. We note that sonication conditions used for the NHEJ assays did not alter the NHEJ viability of WT cells (data not shown).

### Measurement of DNA repair kinetics

50 mL of yeast cultures were harvested at various time points and snap-frozen on dry ice as described above. DNA was extracted by using MasterPure Yeast DNA purification kit (Biosearch Technologies, cat # MPY80200). The DNA samples were quantified and diluted to achieve the same concentration across samples. Primer pairs targeting the DNA break loci and control loci were used to set up PCRs as previousy described^64,74^. The PCR products were then ran on agarose gels, visualized by using Amersham ImageQuant 800 and quantified by using BioRad ImageLab software. The results from 3 independent experiments were averaged and plotted by using Prism 10 (Graphpad).

### Drug Sensitivity (Spot) Assays

Cells were grown in YEP-dextrose overnight to saturation. Next day, they were diluted to a stock concentration of 10^8^ cells/mL. This stock was then serially diluted 10-fold and 2 μL of each dilution was spotted on plates containing hydroxyurea (Sigma-Aldrich, cat # 400046), methyl metanesulfonate (MMS, Sigma-Aldrich, cat # 8207750005), zeocin (Invitrogen, cat # 46-0509) or on YEP-dextrose and exposed to UV with a dose of 200 J/m^2^. Plates were incubated at 30^°^C in the dark until colonies formed, imaged by using Amersham ImageQuant 800 and were processed by using Fiji^118^.

### Yeast RNA extraction, library preparation and RNA sequencing

Cultures of Med12-AID containing cells were grown in YEP-lactate as in the HO-time course experiment. The OD_600_ of the culture was monitored regularly to prevent the cells reaching saturation; cultures were diluted with pre-warmed YEP-lactate as needed to keep them in log phase of growth. Time point samples were harvested as explained above for RNA extraction and snap frozen on dry ice. RNA was extracted by using the RiboPure RNA extraction kit (ThermoFisher, cat # AM1926) and diluted to 100 ng/μL and spiked-in with ERCC spike-in mix (Invitrogen, cat # 4456740) for normalization. The samples were run on 2100 Bioanalyzer (Agilent) to ensure RNA integrity. Sequencing libraries were prepared by using Illumina TruSeq Stranded mRNA Prep and sequenced by using Illumina NovaSEQ X platform (100 bp, paired end). Paired end reads were aligned to SacCer3 reference yeast genome by using STAR^120^. The SAM files were converted to BAM, the reads were sorted and indexed by using Samtools^121^. The read numbers were extracted by using HTseq^122^, and the reads were normalized to ERCC Spike-in controls by using RUVseq^123^. Differential expression analysis was performed by DESeq2^124^. The log_2_ normalized counts were plotted by using Cluster^116^ and Treeview^117^. Aggregate log_2_ fold change values obtained from DESeq2 were plotted by using Prism 10 (Graphpad) and by EnhancedVolcano RStudio package (https://github.com/kevinblighe/EnhancedVolcano).

### Analysis of ChIP-exo data

ChIP-exo raw data was obtained from Rossi et. al. (ref, GEO: GSE147927)^78^. The reads were aligned to SacCer3 reference genome by using bowtie2^125^. The aligned files were sorted and indexed by using Samtools^121^. BAM files were converted to BED by using Bedtools^126^ and then to CPM-normalized bigwig files using bedgraphtobigwig^127^. The normalized bigwig files were plotted using IGV^128^.

### Precision nuclear run-on sequencing (PRO-seq)

PRO-seq was performed as previously published^80^. Briefly, *S. cerevisiae* cells were grown and HO-induced in YEP-lactate as described in detail above. In parallel, *S. pombe* cells, which are used as a spike-in, were grown in YEP-dextrose. The OD_600_ of both yeast cultures were monitored throughout the experiment and cultures were diluted as needed to keep them in log phase of growth. At every time point, 50 million *S. cerevisiae* cells spiked in with 1 million *S. pombe* cells were harvested and permeabilized. To ensure that OD_600_ measurements can accurately predict the cell count and DNA content, especially following the DNA damage, samples collected in parallel were used for DNA extraction. The DNA content of each sample was measured by qPCR to confirm that all collected samples have comparable DNA content (data not shown). Collected samples were permeabilized with 2% sarkosyl (w/v) and snap-frozen in 50 % glycerol (v/v). Sarkosyl-permeabilized cultures were used to set up a nuclear run-on reaction in the presence of biotinylated CTP and UTP (Revvity, cat # NEL542001EA, NEL543001EA). Biotinylated RNA was extracted with phenol-chloroform and purified by using Dynabeads M-280 Streptavidin (Invitrogen, cat # 11206D). 3’ and 5’ ends of the extracted biotinylated RNA were enzymatically ligated with adapter RNA sequences. Following another biotin enrichment of the ligated nascent RNAs, cDNA was prepared by using Superscript III RT (Invitrogen, cat # 18080044). The cDNA was amplified with 2X Phusion (Thermo Scientific) with primers that anneal to the ligated adapter sequences. Libraries were cleaned up and concentrated by ethanol precipitation in the presence of sodium acetate and GlycoBlue (Invitrogen, cat # AM9515). Resulting library samples were sequenced with NovaSEQ X PE 50.

### PRO-seq bioinformatics analysis

The raw PRO-seq reads were cleaned up by using several approaches to improve the alignment rates. First, the adapter sequences were trimmed by cutadapt^129^. Then, low quality and short reads were removed by bbduk (https://sourceforge.net/projects/bbmap). Remaining reads were parsed to remove all the *S. cerevisiae* and *S. pombe* rRNA, tRNA and viral RNA sequences by using bowtie2 unalign function^125^. Remaining reads were aligned to SacCer3^130^ for *S.cerevisiae* or ASM294v3 for *S. pombe*. We note that aligning reads to a catenation of these two yeast genomes did not alter the alignment rates. The number of unique reads that align to spike-in *S. pombe* genome divided by 100000 was used as a scaling factor to normalize the *S. cerevisiae* read counts. To calculate the density of active transcriptional complexes on each gene from *S. cerevisiae* reads, a filtered SAF file was created which excludes overlapping genes, genes with overlapping promoters, ncRNAs, tRNAs, rRNAs, snRNAs, snoRNAs, pseudogenes, transposable elements and genes which are shorter than 400 base pairs; and strand-specific read counts for remaining ∼ 4000 genes were counted using featureCounts^131^. The counts were divided by gene length to calculate RNAPII density per gene and plotted as a histogram by using Prism 10 (Graphpad).

### Chromatin immunoprecipitation

Chromatin immunoprecipitation was performed as previously described^132^ with minor adjustments. The samples from yeast cultures grown in YEP-lactate were collected and cross-linked with a final concentration of 1% formaldehyde, then quenched with glycine at a final concentration of 125 mM. Cross-linked cultures were spun down, washed 3 times with 1 X TBS (pH 7.4) and frozen on dry ice. To lyse the cells, frozen cell pellets were resuspended on ice in 0.55 mL lysis buffer (50 mM HEPES, pH 7.4, 1mM EDTA, pH 8.0, 140 mM NaCl, 1% Triton X-100 (v/v), 0.1 % sodium deoxycholate (w/v)) freshly supplemented with protease inhibitors (1mM ABESF, 0.8 μM aprotinin, 21 μM leupeptin, 15 μM pepstatin, 40 μM Bestatin, 15 μM E64, 600 μM benzamidine), 1 mM PMSF and 1 mg/mL bacitracin. Cell pellets were lysed with 425-600 μm acid-washed glass beads (Sigma-Aldrich, cat # G8772) on a multi-tube vortexer at 4^°^C for 1 hour. Crude cell lysate was harvested and sonicated by using Bioruptor Pico (Diagenode) for 13 cycles (30 seconds on and 30 seconds off) at 4^°^C. Sonicated cell lysates were cleared by centrifugation at maximum speed for 10 minutes at 4^°^C. 50 μL of the crude lysate was set aside as input control, and the remaining cleared lysate was incubated with the antibody of interest (∼1-5 μg) for 1 hour at 4^°^C, and subsequently, with 50-70 μL Protein A or G agarose beads (depending on the antibody) calibrated with lysis buffer for 1 hour at 4^°^C. The antibodies used for the chromatin immunoprecipitation experiments are listed in Table S5. After the antibody incubation, beads were washed twice with lysis buffer, then with lysis buffer with 500 mM NaCl, wash buffer (10 mM Tris-HCl, pH 8.0, 1 mM EDTA, pH 8.0, 0.25 M LiCl, 0.5 % of IGEPAL CA-630 (v/v), 0.5 % of sodium deoxycholate (w/v)) and finally with 1 X TE (10 mM Tris-HCl, pH 8.0, 1 mM EDTA, pH 8.0). Material bound to the beads was eluted with 100 μL of elution buffer (50 mM Tris, pH 8.0, 1 mM EDTA, pH 8.0, 1% SDS) by incubating the beads at 65^°^C for 10 minutes and subsequently with 150 μL of 1 X TE with 0.67 % SDS. Input control samples were supplemented with 200 μL of 1 X TE with 1 % SDS. All samples were treated with 250 μL of 1 X TE and 100 μg of Proteinase K and incubated at 65^°^C overnight for crosslink reversal. The DNA was extracted by using 55 μL of 4 M LiCl and 500 μL of phenol and precipitated by addition of 1 mL of 100 % ethanol and 20 μg of glycogen to the aqueous layer. Samples were incubated at -80^°^C for 1 hour, and DNA was precipitated by centrifugation at maximum speed for 10 minutes at 4^°^C. DNA pellets were washed with 75 % ethanol twice, and after the evaporation of ethanol, were resuspended in 100 μL of 1 X TE. DNA samples were analyzed by using qPCR. Primers used are listed in Table S6.

### ChIP-seq

Sequencing libraries were prepared from γ-H2AX chromatin immunoprecipitation samples by using NEBNext Ultra II DNA library prep kit (NEB, cat # E7645) and NEBNext multiplex oligos for Illumina sequencing (NEB, cat # E7335S, E7500S). Samples were sequenced by using NovaSeq X, PE 100. Reads were aligned by using STAR^120^. RPGC-normalized bigwig files were generated by using bamcompare from the deepTools^133^ kit by ignoring duplicates and mitochondrial chromosome for an effective genome size of 12157105. Matrices for aggregate plots were prepared by using computeMatrix tool from bedTools with a BED file which contains ∼5000 RNAPII-dependent genes, excluding the genes which are not detected by our RNA sequencing analysis. The deepTools plotHeatmap was used with scale-regions command to plot TSS to TES plots, and reference-point command to plot the enrichment around TSS. In addition, RPGC-normalized bigwig files were visualized by using IGV^128^.

### Live cell imaging

Overnight yeast cultures were grown in YEP-lactate at 30°C. To induce DNA damage, cultures were treated with a final concentration of 2 % galactose (w/v) for 8 hours to allow full activation of DNA damage response. Prior to imaging, the cultures were washed into synthetic complete media 3 times and resuspended in synthetic media. Flo11::mNeonGreen Hug1::mScarletI strains were imaged by using Olympus DSU spinning disk confocal microscope with a 100 X 1.45 PlanApo lens in widefield mode. Ddc2::mNeonGreen strains were imaged with Leica SP5 with HC PL APO CS2 63x/1.40 oil lens in z stack mode with a step size of 0.3 μM). The images were processed by using FiJi^118^, while maintaining the same exposure across samples.

### Coimmunoprecipitation

Coimmunoprecipitation was performed as previously described^10^ with minor changes. Briefly, frozen cell pellets were resuspended in lysis buffer (50 mM HEPES-KOH pH 7.5, 150 mM NaCl, 2 mM EDTA, 0.5% IGEPAL CA-630 (v/v)) supplemented with protease inhibitors. After the addition of 200 μL acid-washed glass beads to the resuspension, tubes were vortexed for 60 seconds with MP FastPrep-24 (MP Biomedicals) and cooled on ice for 5 minutes. This bead-beating step was repeated 10 times and percentage of lysis was monitored by checking for “ghost” cells under the light microscope. The extracts were spun down at maximum speed for 10 minutes at 4°C. The protein abundance in precleared crude lysates were quantified by using RC-DC Protein Assay kit (BioRad, cat # 5000121) and diluted to 0.5 mg/mL. 500 μL crude lysates were transferred to fresh tubes on ice and were incubated with Protein A or G-agarose beads (depending on the primary antibody) pre-incubated with the antibody of interest overnight at 4^°^C on a rocker platform. Then, the beads were washed 3 times with 1 mL of lysis buffer supplemented with protease inhibitors for 5 minutes. Samples were eluted in 100 μL 6X Laemmli buffer by boiling at 95^°^C for 5 minutes. Beads were spun down, and the eluates were used for western blotting. Antibodies used for coimmunoprecipitation experiments are listed in Table S5.

### Cell culture

HeLa and U2OS DIvA cells were grown in DMEM with high glucose and pyruvate (Gibco, cat # 11995065), supplemented with 1 % Pen-Strep (Gibco, cat # 15070063), 1 % L-Glutamine (Gibco, cat # A2916801) and 10 % FBS (Avantor, cat # MB1300500) at 37°C with 5 % CO_2_. To induce DNA damage in DIvA cells^88^, cells were treated with 300 nM of 4-hydroxytamoxifen (4-OHT, Sigma, cat # H7904). HeLa cells were treated with 10 μM etoposide for 6 hours to induce DNA damage. To inhibit CDK8, cells were treated with 1 μM BI-1347^95^ for 4 to 6 hours.

### Mammalian Sample preparation for Western Blotting

Harvested cell pellets from 10 cm dishes were resuspended in 550 μL ice cold RIPA buffer (150 mM NaCl, 1 % Triton X-100 (v/v), 0.5 % sodium deoxycholate (w/v), 0.1 % SDS (w/v) and 50 mM Tris-HCl, pH 8.0) supplemented with 1X Halt protease and phosphatase inhibitor (Thermo Scientific, cat # 78440) and incubated on a rotator for 30 mins at 4^°^C. The lysate was cleared by centrifugation at maximum speed for 15 minutes at 4^°^C. The protein concentration in the cleared crude lysate was quantified by using RC-DC Protein Assay (BioRad) and diluted to equal concentrations, mixed with Laemmli loading buffer supplemented with 2-mercaptoethanol. Samples were resolved with SDS-PAGE and blotted as detailed above.

### Nuclear Fractionation

Three 10 cm confluent dishes per condition were harvested by centrifugation and washed with 1 X PBS and combined in one tube. The cell pellets were gently resuspended in hypotonic buffer (20 mM Tris-HCl, pH 7.4, 10 mM KCl, 2 mM MgCl_2_, 1 mM EGTA, 340 mM sucrose, 10 % glycerol (v/v)) supplemented fresh with 1 X Halt phosphatase and protease inhibitor and 0.5 mM DTT. After 5 minutes of incubation on ice, Triton X-100 was added to a final concentration of 0.1 % (v/v) into each tube and samples were mixed by gentle inversion. Samples were spun down at 1000 rcf for 5 minutes at 4^°^C to separate the cytoplasmic fractions (supernatant) and intact nuclei (pellet). Pellets containing the nuclei were washed with isotonic buffer (20 mM Tris-HCl, pH 7.4, 150 mM KCl, 2 mM MgCl_2_, 1 mM EGTA, 340 mM sucrose, 10 % glycerol (v/v)), distributed equally into three tubes (for nucleoplasmic extraction with three different salt concentrations) and spun again. Supernatants were discarded, each cleared nuclear pellet was treated with buffers with increasing salt concentrations: buffer 1 containing 150 mM KCl, buffer 2 containing 250 mM KCl and buffer 3 containing 400 mM KCl, supplemented with 1 X Halt phosphatase and protease inhibitor and 0.5 mM DTT. Nuclei were incubated on ice for 30 minutes to salt-extract the chromatin-bound proteins. After the incubation, samples were spun down at 2000 rcf for 5 minutes. Supernatants were transferred into clean tubes as nucleoplasmic fractions. Pellets, which contain the chromatin-bound proteins and DNA, were roughly resuspended in 1 X PBS supplemented with CaCl_2_ and solubilized with benzonase treatment of the DNA on ice. After the incubation, samples were further solubilized with RIPA buffer. A final concentration of 2X Laemmli buffer was added to all samples. Samples were blotted for Histone H3 (to probe for chromatin fractions), SP1 (to probe for nucleoplasmic fractions) and GAPDH (to probe cytoplasmic fractions). The antibodies used for these experiments are listed in Table S5.

### CRISPR/Cas9 gene targeting in HeLa

1 μM of sgRNA (IDT), which contains the gRNA sequence CGAGGACCTGTTTGAATACG targeting the CDK8 locus, was incubated 1 μM Cas9-RFP (IDT), Cas9 Plus (from Lipofectamine CRISPRMAX kit, Invitrogen, cat # CMAX00001) and Opti-MEM in a final volume of 25 μL for 5 minutes at room temperature to form an RNP complex as per manufacturer’s instructions. RNP complex was then mixed with 1.2 μL of CRISPRMAX transfection reagent from Lipofectamine CRISPRMAX kit and 25 μl of Opti-MEM and incubated at room temperature for 20 minutes to form a transfection complex. Cells grown in DMEM-FBS without antibiotics were trypsinized, diluted and transfected with the transfection mix. One day after the transfection, cells were trypsinized and sorted into 96-well plates with FACSAria Fusion 5-18 (BD Biosciences) on the RFP channel. Single cells were expanded and then genotyped with PCR after extracting the DNA with QuickExtract DNA extraction solution (Biosearch Technologies, cat # QE0905T). Two CDK8 knockout clones were selected and further confirmed with western blotting and premium PCR sequencing with primers flanking the editing site (Plasmidsaurus), then tested for mycoplasma contamination with PCR and microscopy-based methods as detailed above. Single wild-type HeLa cells were sorted and expanded in parallel as a control for the downstream experiments, to rule out any possible convoluting effects of clonal selection. The primer sequences used for CDK8 knockout genotyping are listed on Table S6.

### DsiRNA for CDK8 knockdown

HeLa cells were plated to confluency 24 hours prior to transfection. Transfection mix was prepared by using Lipofectamine 2000 (Invitrogen, cat # 11668027) and Opti-MEM as per manufacturer’s recommendations with a duplex DsiRNA sequence targeting CDK8 (AGGCAUUAUACCAAAGCUAUUGATA) and a control DsiRNA sequence (IDT) at a final concentration of 2 nM. Transfected cells were expanded and harvested 48 to 72 hours after knockdown. The efficiency of knockdown was determined with a RT-q-PCR and Western blotting. The primer sequences used for CDK8 qPCR are listed in Table S6.

### Immunofluorescence

DIvA cells were seeded on coverslips at 50 - 70 % confluency and grown overnight to attach. 4 hours after treatment with 4-hydroxytamoxifen, cells were washed twice with 1 X PBS and fixed by incubation in ice cold fixative solution (3 % formaldehyde (w/v), 3.4 % sucrose (w/v) in 1 X PBS) for 10 minutes. Fixative solution was removed, and the coverslips were washed 3 times with 1 X PBS to remove residual formaldehyde. Cells were permeabilized by incubation in freshly made permeabilization buffer (20 mM HEPES-NaOH, pH 7.4, 50 mM NaCl, 3 mM MgCl_2_, 300 mM sucrose, 0.5 % Triton X-100 (v/v)) for 10 minutes. Then, they were incubated in blocking buffer (1 % BSA (w/v) in 1 X PBS) overnight at 4^°^C. The next day, the coverslips were incubated in 1:500 anti-γ-H2AX antibody coupled with AlexaFluor-555 (Millipore, cat number # JBW30) in blocking buffer for one hour at room temperature. The coverslips were washed 3 times with 1 X PBS, air-dried and mounted with ProLong Diamond Antifade with DAPI (Invitrogen cat # P36971). The images were taken with Leica SP5 with HC PL APO CS2 63x/1.40 oil lens and processed using FiJi^118^. The foci number and intensity were quantified by using CellProfiler^134^. More than 700 foci from 50 cells from two independent staining experiments per condition were analyzed. One-way ANOVA was used for the statistical analysis of foci number. Unpaired t-test was used for the statistical analysis of foci intensity (*** *p*<0.0001).

### Clonogenic survival assay

Cells were plated on 12-well plates at a density of 100 - 200 cells/well and grown for 12 hours to attach. Then, they were treated with various concentrations of etoposide (Sigma Aldrich, cat # E1383), topotecan (TPT, Sigma Aldrich, cat # 1672257), zeocin and hydroxyurea (Sigma Aldrich, cat # 400046). 24 hours after the treatment, the drug containing spent media was removed and exchanged with pre-warmed fresh media. 10 to 14 days after plating, when the colony sizes reach ∼ 50-100 cells, the wells were washed with 1X PBS. Then, the cells were incubated in crystal violet fixing and staining solution (20 % methanol (v/v), 0.5 % of crystal violet (w/v)) for 5 minutes. After the incubation, the plates were washed with distilled water, air-dried and imaged with G:BOX F3 (Syngene) on a backlit light plate. The images were cropped with FiJi^118^ and colony numbers were quantified by using CellProfiler^134^. The colony numbers in the treated plates were normalized to untreated controls and plotted using Prism 10 (Graphpad).

## Notes

### Summary of Updates

We made substantial changes in the updated version of the paper: - Figure 1 and 2 are merged. - Updated Figure 2 contains additional drug sensitivity assays. - RNAseq assay shown in the old Figure 4 was redone with three replicates. We also performed additional analyses with this dataset, comparing it to previously published DNA damage transcriptome datasets (which is included in the supplement). - In addition to bulk RNAseq, we performed PROseq to assay nascent transcription following DNA damage in WT and cdk8-kd cells. - To show that transcriptional output modulates DNA damage response, we employed several RNAPII mutants and analyzed their responses to DNA damage. - We analyzed changes to Cdk8 kinase activity upon DNA damage via Med2 phosphorylation and Flo11 expression. - We repeated the coimmunoprecipitation assays shown in the old version Figure 5 in cdk8-as mutants. To support these assays, we introduced mutations in CKM-core Med binding interface and assayed the DNA damage response of these mutants. - In addition to the ChIP assays shown in the old version of Figure 6, we performed ChIP seq to analyze the genome-wide distribution of g-H2AX in the absence of Cdk8 kinase activity. - We expanded our investigation of the CKM effect on DNA damage response to human cell lines. For this, we employed knockout and knockdown human lines to show that DNA damage leads to RNAPII eviction in a CDK8-dependent manner. - We studied the DNA damage response and sensitivity of CDK8 knockout lines. - We examined the DNA damage foci in human lines following CDK8 inhibition. - Lastly, the paper now contains accompanying supplemental figures.

